# Description of *Pectobacterium pacificum* sp. nov. isolated from *Colocasia esculenta*

**DOI:** 10.1101/2022.02.08.479620

**Authors:** Mohammad Arif, Komal, Diksha Klair, Dario Arizala, Aarya Mishra, Gamze Boluk, Małgorzata Waleron, Anne M. Alvarez, Shefali Dobhal

## Abstract

*Pectobacterium* comprises Gram-negative, pectinolytic plant pathogens that cause soft rot and blackleg diseases on numerous economically important crops. During sampling in 2018 in taro (*Colocasia esculenta*) fields in Hawai’i, two bacterial strains PL152^T^ and PL155 were isolated from taro. Peliminary molecular characterization placed both strains in the genus *Pectobacterium*. However, their taxanomic postions could not be resoved at the species level. A polyphasic taxonomic approach incorporating phenotypic characterization, 16S rRNA gene sequence analysis, multilocus sequence analysis (MLSA), whole-genome-based phylogeny, average nucleotide identity (ANI), and digital DNA–DNA hybridization (dDDH) was therefore employed to determine their taxonomic status. Phylogenetic and phylogenomic analyses consistently placed PL152□ and PL155 together in a distinct and well-supported lineage within the genus *Pectobacterium*, separate from currently recognized species. The two strains exhibited high genomic relatedness to each other, whereas the average nucleotide identity (ANI) and digital DNA-DNA hybridization (dDDH) values, were lower than the accepted species delineation thresholds of 95% and 70%, respectively, supported the delineation of a novel bacterial species. The genomic characteristic of PL152 included a genome size of 5.02 Mb and a DNA G+C content of 52%. Collectively, based on a polyphasic approach including phylogenetic, phlyogenomic, biochemical, and phenotypic evidences supports the classification of strains PL152□ and PL155 as representatives of a novel species within the genus *Pectobacterium*, for which the name *Pectobacterium pacificum* sp. nov. is proposed. Strain PL152 (= ICMP 24362□; LMG 32536□) is designated as the type strain.

## INTRODUCTION

The genus *Pectobacterium*, previously classified within the genus *Erwinia*, is a significant group within the family *Pectobacteriaceae* (Adeolu et al., 2016). These bacteria are characterized as Gram-negative, fermentative, rod-shaped phytopathogens primarily known for their ability to degrade plant tissues. Their pathogenicity arises from the secretion of diverse plant cell wall degrading enzymes (PCWDEs), such as cellulases, pectinases, and proteases, which effectively break down plant structural polysaccharides and proteins. These enzymatic activities lead to maceration of plant cell walls and tissues, creating an optimal environment for bacterial colonization, promoting nutrient release, and facilitating infection and symptom development (Arif et al, 2022; Arizala and Arif, 2019; Hugouvieux-Cotte-Pattat et al., 2014). *Pectobacterium* species are recognized globally as among the ten most economically important and extensively studied plant pathogens (Mansfield et al., 2012). This prominence is attributed to their broad host range, infecting numerous agriculturally important crops and causing substantial economic losses through diseases such as soft rot, blackleg, and wilt (Mansfield et al., 2012; Klair et al, 2021; Toth et al, 2021). The symptoms of these diseases can vary significantly, ranging from tissue maceration and discoloration to complete plant collapse, severely impacting crop yield and quality (Czajkowski et al., 2011; Klair et al, 2021, 22). The versatility of *Pectobacterium* extends beyond plant infection, as species within this genus have been isolated from diverse ecological niches, including soil, groundwater, and various invertebrate hosts (Toth et al, 2021). These alternative reservoirs potentially aid in the dissemination and persistence of *Pectobacterium* in both agricultural and natural ecosystems, emphasizing the complexity of managing these pathogens effectively (Glasner et al., 2008). The ecological complexity of this genus underscores the importance of accurate taxonomic classification and the description of novel species for strains that do not match any existing species because they fail to meet the established average nucleotide identity (ANI) and digital DNA-DNA hybridization (dDDH) thresholds of 95-96% and 70%, respectively.

The taxonomy of the genus *Pectobacterium* has undergone continuous revision and reorganization, reflecting advances in bacterial classification and genomic technologies (Parte et al., 2020; Portier et al., 2020; Oren and Garrity, 2022). Figure 1 illustrates the taxonomic history of the genus, including its establishment, major taxonomic revisions, species transfers, and the rapid expansion in the number of recognized species over time. Next-generation sequencing (NGS), particularly whole-genome sequencing, has significantly advanced these efforts, enabling detailed analysis of taxonomically complex genera and enhancing comparisons of phylogenetic and evolutionary relationships among species and subspecies of plant pathogenic genera (Arif et al., 2022; Zhang et al., 2016). Species within the genus *Pectobacterium* have been recovered from a wide range of plant hosts and environmental sources. The type strains of several *Pectobacterium* species were originally isolated from potato, including *P. carotovorum*, *P. brasiliense*, *P. atrosepticum*, *P. versatile, P. punjabense*, *P. parmentieri*, *P. parvum*, *P. polare (P. polaris)*, *P. peruviense*, and *P. sinaloense*. While the type strains of other *Pectobacterium* species were originally isolated from non-potato hosts, such as *P. actinidiae* from kiwifruit, *P. odoriferum* from *Cichorium intybus*, *P. wasabiae* from wasabi, *P. aroidearum* from aroid/monocot hosts, *P. betavasculorum* from sugar beet, *P. araliae* from Japanese angelica tree, *P. colocasium* from taro, *P. jejeuense* from cucumber, *P. zantedeschiae* from calla lily, and *P. cacticida* from cactus (recently been proposed for reclassification as *Alcorniella cacticida* comb. nov., however, the proposed name has not been validly published. In contrast, species such as *P. aquaticum*, *P. fontis*, *P. polonicum*, and *P. quasiaquaticum* were originally recovered from aquatic or groundwater sources. This broad distribution highlights the ecological diversity of *Pectobacterium* species across crop hosts, ornamental plants, and environmental reservoirs. Currently, the genus comprises more than twenty validly published species recovered from diverse plant hosts and environmental reservoirs, reflecting the remarkable ecological and taxonomic diversity within two proposed species, *P. colocasium* (Zhou et al., 2022) and *P. zantedeschiae* (Waleron et al., 2019a), as well as the proposed reclassification of *P. cacticida* as *Alcorniella cacticida* comb. nov. (Jonca et al., 2024), which have not yet been validly published under the International Code of Nomenclature of Prokaryotes (ICNP).

**Figure 1.**
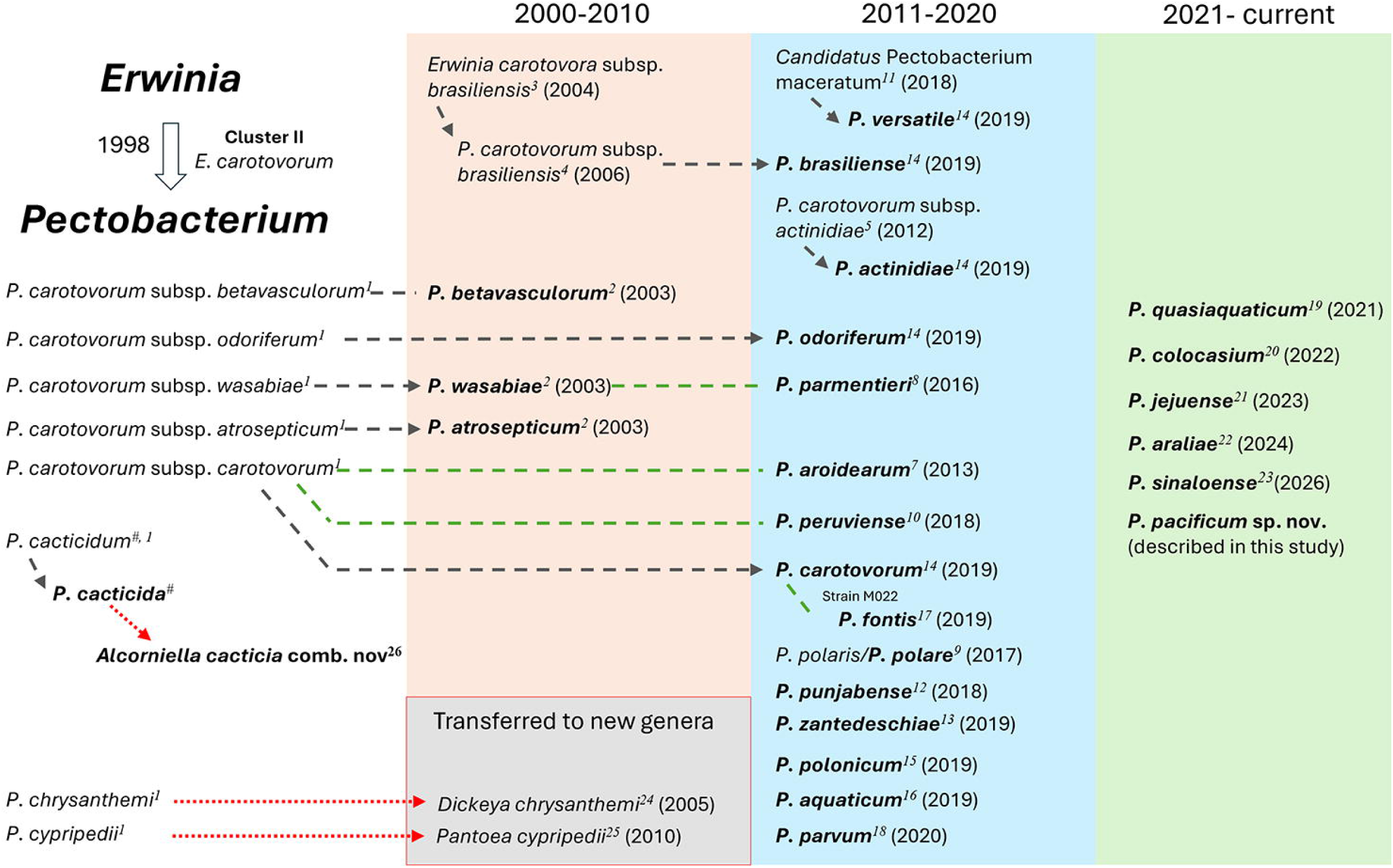
Historical overview of taxonomic changes within the genus *Pectobacterium.* The timeline illustrates key milestones in the taxonomy of *Pectobacterium*, including the original description of species, recognition of subspecies, elevation of subspecies to species, proposal of new species, and major taxonomic revisions. Black dashes indicate either elevation to species level or change of name. Green dashes indicate that some strains moved to new species. Red dashes indicates that the species transfer to new genera. Names with bold letters indicate the current name of the species as of August 2026. ^1^Hauben et al., 1998; ^2^Gardan, et al., 2003; ^3^Duarte, et al., 2004; ^4^El Tassa and Duarte, 2006 (Use of the name *Pectobacterium carotovorum* subsp. *brasiliensis* by El Tassa and Duarte; not validly published); ^5^Koh et al., 2012; ^6^Nabhan et al., 2012; ^7^Nabhan, et al., 2013; ^8^Khayi, et al. 2016; ^9^Dees, et al. 2017; ^10^Waleron, et al. 2018; ^11^Shirshikov, et al., 2018; ^12^Sarfraz, et al., 2018; ^13^Waleron et al., 2019; ^14^ Portier et al., 2019; ^15^Waleron et al., 2019; ^16^Pedron et al., 2019; ^17^Oulgazi et al., 2019; ^18^Pasanen, et al., 2020; ^19^ Ben Moussa et al., 2021; ^20^Zhou et al., 2022; ^21^Hong et al 2023; ^22^Sawada et al., 2024; ^23^Valdez-López, et al., 2026; ^24*^Samson et al., 2005 (the genus *Dickeya* was proposed and transferred *Pectobacterium chrysanthemi* to this genus as *Dickeya chrysanthemi* comb. nov.); ^25*^Brady et al., 2010 (Brady et al. later transferred this species to the genus *Pantoea* as *Pantoea cypripedii* comb. nov.); ^#^Alcorn et al., 1991 (*Erwinia cacticida* sp. nov. Hauben et al., 1998 transferred it to *Pectobacterium cacticidum* which in 1999 was corrected to *Pectobacterium cacticida* under Rule 61, Validation List No. 68), ^26^Jonca et al., 2024 (proposal for reclassification *Pectobacterium cacticida* as *Alcorniella cacticida* comb. *nov.*, not approved yet)

In this study, two strains, PL152^T^ (ICMP 24362^T^; LMG 32536^T^), and PL155 isolated from taro, were analyzed and proposed as a novel species, *Pectobacterium pacificum* sp. nov. This research aims to clarify the taxonomic position of these strains through comprehensive genomic and phenotypic characterization.

## MATERIALS AND METHODS

### Bacterial strain, isolation and identification

The strains PL152^T^ and PL155, referred to hereafter as *P. pacificum*, were isolated from taro corm samples exhibiting soft rot symptoms collected in Oahu, Hawaii in 2018. Infected taro corms were surface sterilized using a 0.6% sodium hypochlorite solution for 30 seconds, followed by three consecutive rinses with distilled water, each for 30 seconds. The corm tissues were then aseptically macerated in sterile 1.5 ml eppendorf tubes using sterile pestles. The resulting tissue suspension was streaked onto Crystal Violet Pectate (CVP) medium (Hélias et al., 2012), allowing for colony isolation based on the formation of characteristic cavities due to pectate degradation. Colonies producing pits after 24 hours of incubation were purified through two consecutive rounds of re-streaking on nutrient agar supplemented with 0.4% dextrose (NAG). The identity of the purified bacterial colonies was confirmed using PCR assays with *Pectobacterium*-specific primers (Arizala et al., 2022). Finally, pure cultures derived from single colonies were preserved at -80°C in a solution consisting of 30% glycerol mixed with LB liquid medium (30:70 v/v). For identification confirmation of bacterial strains PL152^T^ and PL155, the bacterial DNA was isolated from purified colonies using the DNeasy Blood and Tissue kit following the manufacturer’s protocol (Qiagen, Valencia, CA). Three housekeeping genes, chromosomal replication initiator protein DnaA (*dnaA*), glyceraldehyde-3-phosphate dehydrogenase (*gapA*), and DNA topoisomerase ATP-hydrolyzing subunit B (*gyrB*), were used for identification. All genes were amplified using the previously reported primers and conditions (Boluk et al., 2020; Dobhal et al., 2020). The amplified PCR products were enzymatically cleaned using ExoSAP-IT (Affymetrix, Santa Clara, CA) sequenced at Genewiz facility (La Jolla, CA) using both forward and reverse primers, and further aligned and manually edited to remove errors using Geneious R10 (Kearse et al., 2012). The consensus sequences of *dnaA*, *gapA* and *gyrB* gene sequences for PL152^T^ and PL155 strains were submitted to the NCBI GenBank database under the accession numbers: PL152-MZ542535 (*dnaA*), MZ542537 (*gyrB*), MZ542539 (*gapA*) and for PL155-MZ542536 (*dnaA*), MZ542538 (*gyrB*), and MZ542540 (*gapA*).

### Genome sequencing, assembly, and overall genome relatedness indices

For whole-genome sequencing, bacteria were recovered from stored pure cultures of strains PL152^T^ and PL155, and streaked onto nutrient agar supplemented with 0.4% dextrose for genomic DNA isolation. A half-loopful of pure bacterial culture was used for DNA isolation with a QIAGEN Genomic-tip 100/G (Qiagen). The DNA was quantified using a Qubit 4 fluorometer with Qubit dsDNA HS kit (Thermo Fisher Scientific, Waltham, MA), and analyzed on a 1.5% agarose gel. The genomes of PL152^T^ and PL155 were sequenced using Oxford Nanopore and Illumina sequencing platforms. For Oxford Nanopore sequencing, the genomic library of strain PL152 was prepared using the Native Barcoding Expansion Kits (EXP-NBD104 and EXP-NBD114) in combination with the Ligation Sequencing Kit (SQK-LSK109).

In contrast, the genomic library of strain PL155 was prepared using the Native Barcoding Kit 24 V14 (SQK-NBD114.24). A total of 400 ng of high-quality genomic DNA was used for library preparation for each strain. Genome sequencing was performed on a MinION device using an FLO-MIN112 (R10.4.1) flow cell according to the manufacturer’s instructions (Oxford Nanopore Technologies, Oxford, UK). Sequencing data generated with the flow-cell was monitored in real time using the MinKNOW software version 4.0.20 (Oxford Nanopore Technologies, Oxford, UK). The obtained FAST5 sequences from MinION were base-called using MinKNOW version 4.0.20 (Oxford Nanopore Technologies, Oxford, UK). For Illumina sequencing, the DNA libraries were prepared using Seqwell plexWell LP384 Library Preparation kit (seqWell, Beverly, MA) using 10 ng gDNA. The prepared libraries were amplified with 8 PCR cycles, analyzed using Bioanalyzer 2100 (Agilent, Santa Clara, CA), quantified with Qubit, and combined into two pools at equimolar ratios. The library pool was quantified by qPCR with a Kapa Library-Quant kit (Kapa Biosystems/Roche, Basel, Switzerland) and sequenced using an Illumina NovaSeq system (Illumina, San Diego, CA) with paired-end 150-bp reads. The Illumina library preparation and sequencing was performed at Novogene facility (Sacramento, CA).

*De novo* genome assemblies were generated using Oxford Nanopore long reads together with paired-end (2 × 150 bp) Illumina short reads. First, the genomes of strains PL152□ and PL155 were assembled in CLC Genomics Workbench version 20.0.1 (Qiagen, Denmark) using the “Long Read Support” tool with the “*De novo* Assembly Long Reads” option. The resulting long-read assemblies were subsequently polished with the Illumina short reads using the “Polish with Reads” tool. The completeness and contamination of both genome assemblies were assessed using CheckM version 1.2.3 (Parks et al., 2015), specifying the genus *Pectobacterium* as the taxonomic marker. The generated complete genomes were annotated using the Prokaryotic Genome Annotation Pipeline (PGAP) version 4.8 (Tatusova et al., 2016). To support the designation of strains PL152□ and PL155 as representatives of a novel species, *P. pacificum*, comprehensive genome-wide similarity analyses were conducted. Average Nucleotide Identity (ANI) values were calculated using the CLC Genomics Workbench, while digital DNA-DNA hybridization (dDDH) values were determined using the Genome-to-Genome Distance Calculator (GGDC, version 3.0), employing the recommended formula 2.

### Phylogeny based on 16S rRNA, multi-locus sequence analysis and whole genome

For 16S rRNA phylogenetic analysis, the 16S rRNA gene sequences of strains PL152□ and PL155 were extracted from the whole-genome sequences and submitted to GenBank. To confirm the generic identification of strains PL152□ and PL155, comparative 16S rRNA gene sequence analyses were performed using the Type (Strain) Genome Server (TYGS; https://tygs.dsmz.de), a platform for genome-based taxonomic and phylogenomic assessment of prokaryotes. For this analysis, the TYGS server was configured in a restricted mode, allowing comparisons only among the type strains provided by the user, and excluding type strains available in the public TYGS database. The type strains of all *Pectobacterium* species *P. colocasium* LJ1^T^, *P. actinidiae* KKH3^T^, *P. carotovorum* ICMP5702^T^, *P. aroidearum* L6^T^, *P. brasiliense* LMG21371^T^, *P. polaris* NIBIO1006^T^, *P. parvum* s0421^T^, *P. odoriferum* NCPPB3839^T^, *P. versatile* CFBP6051^T^, *P. aquaticum* A212-S19-A16^T^, *P. zantedeschiae* 9M, *P. fontis* M022^T^, *P. polonicum* DPMP315^T^, *P. punjabense* SS95^T^, *P. parmentieri* RNS08-42-1A^T^, *P. wasabiae* CFBP3304^T^, *P. atrosepticum* 21A^T^, *P. peruviense* IFB5232^T^, *P. betavasculorum* NCPPB2795^T^, *P. quasiaquaticum* A477-S1-J17^T^, *P. cacticida* CFBP3628^T^, *P. jejuense* 13-115^T^ *P. araliae* MAFF 302110 ^T^ and *P. sinaloense* LFLA-215^T^, along with the whole genomes of PL152^T^ and PL155, were uploaded to the TYGS server (Supplementary Table 1). The 16S rRNA phylogenetic tree generated using the TYGS was visualized using the web based tool interative Tree of Life (Itol V7, https://itol.embl.de).

For taxonomic classification and novel species delineation, multi-locus sequence analysis (MLSA) (De Vos, 2011) was performed using nine housekeeping genes: nucleotide sequences of *dnaA* (chromosomal replication protein), *gapA* (glyceraldehyde-3-phosphate dehydrogenase), *gyrB* (DNA gyrase subunit B), *recA* (recombinase A), *dnaN* (DNA polymerase III subunit beta), *rpoS* (RNA polymerase sigma factor RpoS), *mdh* (malate dehydrogenase), *rpoA* (DNA-directed RNA polymerase subunit alpha), and *dnaK* (molecular chaperone DnaK) were retrieved from annotated genome assemblies from the NCBI Genome database for 57 *Pectobacterium* strains, including the type strains, together with strains PL152□ and PL155. The strain and accession numbers are provided in Supplementary Table 2. Individual gene sequences were extracted from the annotated genome assemblies. The sequences were aligned using ClustalW, trimmed, and concatenated using Geneious Prime version 2025.0.3 (Kearse et al., 2012). The final concatenated alignment used to construct the phylogenetic tree comprised of 9,308 bp. Phylogenetic relationships among the strains were inferred using a maximum-likelihood (ML) approach implemented in RAxML v8.2.11 within Geneious Prime, using the GTR+GAMMA nucleotide substitution model, and branch support was assessed using 1,000 rapid bootstrap replicates. Whole-genome phylogenetic analysis was conducted using strains PL152 and PL155 together with the type strains within the genus *Pectobacterium*. The genomes of all *Pectobacterium* type strains were retrieved from the NCBI database and uploaded to the Type (Strain) Genome Server (TYGS). The Newick file generated by Geneious and TYGS were uploaded to the Interactive Tree of Life (iTOL) version 7 for visualization and annotation of the phylogenetic tree. The taxonomic status of these strains was further assessed using the Type Strain Genome Server TYGS; https://tygs.dsmz.de/), an automated platform for genome-based prokaryotic taxonomy. TYGS compares submitted genomes against a curated and regularly updated database of bacterial type-strain genomes and uses digital DNA-DNA hybridization (dDDH) values, including the accepted 70% species delineation threshold, to identify potential novel species (Meier-Kolthoff and Goker, 2019; Meier-Kolthoff et al., 2022).

### Genomic Analysis

The *Pectobacterium* genome sequences were analyzed to identify genes involved in chemotaxis and flagellar biosynthesis. Genome annotations were used to identify genes encoding flagellar proteins, chemotaxis proteins, and methyl-accepting chemotaxis proteins (MCPs). The genetic searches were carried out using Geneious Prime using Map to Reference function. The identified genes were grouped by function and compared among genomes to assess their presence and possible role in motility, host colonization, and disease development. Furthermore, the genomes were analyzed for the presence of genes encoding PCWDEs based on functional annotations, with emphasis on enzymes commonly associated with soft rot development, including pectate lyases, polygalacturonases, cellulases, xylanases, proteases, and other carbohydrate-active enzymes. In addition, genes associated with bacterial secretion systems were identified to assess potential mechanisms involved in host interactions, enzyme delivery, and pathogenicity. This analysis was used to support the genomic characterization of the proposed novel species and to better understand its potential pathogenic features.

The pan-genome analysis was performed using Panaroo to construct polished pan-genome, generate presence and absence matrix and to identify unique gene clusters (https://github.com/gtonkinhill/panaroo). The annotated genomes files of PL152^T^ and PL155 and a subset of closely related genomes, were analyzed (Tonkin-Hill et al., 2020). These genes were then grouped into functional categories based on their annotations. The presence of microbial resistance was perfomed using AMRFinderPlus (Feldgarden et al., 2021). Biosynthetic gene clusters related to secondary metabolite production, including potential phytotoxins, antibiotics, and antimicrobial compounds, were predicted using antiSMASH version 8.0.4 (Blin et al., 2021). Putative anti-phage defense system were identified using DefenseFinder v2.0.0 (Tesson et al., 2022; Neron et al., 2023). BAGEL4 (<u>BAGEL4 - RiPPs miner</u>) was used to screen both genomes for antimicrobial compunds (Van Heel et al., 2018). Additionally, Clustered Regularly Interspaced Short Palindromic Repeats (CRISPRs) arrays and the type of CRISPR associated proteins (Cas) systems were predicted using CRISPRCASFinder (<u>CRISPR-CAS++</u>) in both genomes PL152^T^ and PL155 (Couvin et al., 2018).

### Phenotypic, biochemical characterization, pathogenecity and antibiotic sensitivity assay

Strains PL152^T^ and PL155 were evaluated for swimming and swarming motility using semi-solid agar media as previously described (Chen et al, 2016). Briefly, the swimming medium consisted of 10 g/L tryptone, 5 g/L NaCl, and 3 g/L agar, whereas the swarming assay medium contained 10 g/L tryptone, 5 g/L NaCl, and 4 g/L agar. Each medium was inoculated by spotting 5 µL of an overnight bacterial culture adjusted to an optical density (OD) of 1.0 at 600 nm onto the center of the plate. The plates were incubated at 28, after which the diameter of bacterial growth was measured and recorded. Each experiment was conducted in duplicate. Biochemical characterization of the novel strains PL152 and PL155 was performed using the Biolog GEN III MicroPlate system (Biolog, Inc., Hayward, CA). This system evaluates bacterial metabolism through 71 carbon source utilization assays and 23 chemical sensitivity assays. Pure bacterial cultures were streaked onto BUG agar (Biolog, Inc.) and incubated at approximately 28 °C for 24 h. Individual colonies were then selected with sterile loops and suspended in inoculating fluid B to prepare a standardized inoculum. Subsequently, 100 µL of the bacterial suspension was dispensed into each well of a 96-well GEN III MicroPlate. The plates were incubated at 28 °C for 24 h, and the results were recorded using a Biolog MicroStation ELx808BLG Reader (Biolog). The results were recorded as positive or negative for carbon source utilization and as sensitive or resistant for the chemical sensitivity assays.

Biochemical properties of the novel strains PL152 and PL155 were investigated using the API 20E and API ZYM biochemical test kits (bioMérieux, Marcy-l’Étoile, France). These tests were used to assess biochemical reactions and enzymatic activities. Bacterial cultures were grown on NAG medium, and colonies from 24-hour-old plates were used for testing. For the API 20E strips, comprising 21 miniaturized biochemical tests, bacterial inocula were prepared by suspending a single isolated colony in API suspension medium (0.85% NaCl). The suspension was immediately dispensed into the test tubes and cupules according to the manufacturer’s instructions. Tests were incubated at 28±2°C, and results were recorded after 24 hours. Enzymatic activities of the bacterial strains were analyzed using the API ZYM assay, which semi-quantitatively identifies 19 different enzymatic reactions. To prepare the inoculum, bacterial colonies were suspended in API suspension medium, adjusting the turbidity to match the 5 McFarland. A volume of 65 µL of this prepared suspension was then dispensed into each cupule of the API ZYM test strip. The inoculated strips were incubated at 28±2°C for 4 hours and 30 minutes. After incubation, enzymatic reactions were visualized by adding one drop each of ZYM A and ZYM B reagents to each cupule. Results were determined by observing color changes approximately 5 minutes after reagent addition, indicating enzymatic activity. Cytochrome oxidase activity was assessed using OxiStrips (Hardy Diagnostics, Santa Maria, CA); a positive reaction was indicated by a dark blue color change within 90 seconds. Catalase activity was verified by adding 3% hydrogen peroxide (H□ O□), with bubble formation signifying a positive result. The production of extracellular enzymes, including pectate lyase (pel) and cellulase (cel), was evaluated using a semi-quantitative assay as described previously (Chatterjee et al, 1995). For the assays, 50 µl of bacterial suspension (OD□ □ □ ≈ 1.0) was dispensed into wells on media plates and incubated at 28°C. Pectate lyase (pel) activity was visualized after 10 hours by flooding the plates with 4N HCl, while cellulase (cel) activity was detected using 2% Congo red followed by 5 M NaCl treatment. Pectinolytic activity was also evaluated by spot inoculation on CVP medium, with pit formation monitored after incubation at 28°C. The assay was performed in duplicates (Dobhal et al., 2025).

The ability of the bacterial strains to cause tissue maceration was assessed through pathogenecity assays conducted on both potato tubers and taro corms following the protocol described by Dobhal et al (2025). Briefly, to ensure surface sterility, each tuber or corm was treated with 0. 6% sodium hypochlorite for 10 minutes, followed by three consecutive rinses with sterile distilled water, each lasting 3 minutes and aseptically sliced using a sterile knife. Each slice was carefully inoculated with 20 µl of bacterial suspension (OD□ □ □ ≈ 1.0) and incubated at ambient room temperature (approximately 26°C). Symptoms of tissue softening or rot were observed and recorded after 24 hours. Each assay was performed in triplicate to ensure reproducibility of the results.

Both strains, PL152□ and PL155, were tested for antibiotic sensitivity using the disc diffusion method. A 100 µl aliquot of bacterial suspension (OD□ □ □ ≈ 1.0) was uniformly spread onto nutrient agar medium supplemented with 0.4% dextrose. Sterile filter paper discs were each absorbed with 20 µl of different antibiotic solutions: penicillin (50 mg/ml), kanamycin (50 mg/ml), tetracycline (40 mg/ml), chloramphenicol (50 mg/ml), carbenicillin (100 mg/ml), Gentamicin (50 mg/ml), and bacitracin (50 mg/ml). The discs were then placed at the center of the inoculated plates, which were subsequently incubated at approximately 28°C for 24 hours.

### Transmission Electron Microscopy (TEM)

TEM was used to observe bacterial morphology. Sample preparation followed the protocol provided by the Biological Electron Microscope Facility at the University of Hawai i at Mānoa and as described by Dobhal *et al*. (2025). Briefly, bacterial cultures were grown overnight in nutrient broth at 28°C with shaking at 120 rpm. For imaging, cells were placed on glow-discharged, 200-mesh, carbon/Formvar-coated copper grids, then negatively stained with 1% uranyl acetate and rinsed three times with sterile distilled water. The grids were examined using a Hitachi HT7700 TEM operated at 100 kV. High-resolution images were acquired with an AMT BioSprint16M-ActiveVu CCD camera (16 megapixels, 4895 × 3920 resolution). Bacterial cell length and diameter were measured in triplicate.

## RESULTS AND DISCUSSION

### Genomic Features and Overall Genomic Relatedness Indices (OGRIs)

The assembled genomes of strain PL152□ and PL155 were submitted to the NCBI GenBank genome database under the accession number CP091064.2 and CP118921.2, respectively. The genome sizes were 5,022,655 bp and 5,024,691 bp, respectively, with no plasmids detected. The G+C content was 52 mol% for both genomes. Results from the NCBI Prokaryotic Genome Annotation Pipeline (PGAP) indicated that the PL152□ genome encodes a total of 4,510 genes, including 4,401 coding sequences (CDSs), 109 RNA genes (including 22 rRNAs and 77 tRNAs), 3 CRISPR arrays, and 161 pseudogenes. Similarly, the PL155 genome encoded 4,524 genes, including 4,415 CDSs, 109 RNA genes (including 22 rRNAs and 77 tRNAs), 3 CRISPR arrays, and 2,26 pseudogenes. Genome quality assesment using CheckM showed 0.49 and 0.37% conatmination for the PL152□ and PL155 genomes (Supplementary Table 3). Comparison and annotation using the RASTtk Model SEED database categorized the genome features of *Pectobacterium pacificum* strains PL152□ and PL155 into 28 distinct functional groups (Brettin et al., 2015). The three largest functional categories for both strains are amino acids and derivatives (PL152□ and PL155-330 and 336 features, respectively), carbohydrates (PL152□ : and PL155: 286 and 314 features, respectively), and protein metabolism (PL152□ and PL155: each with 216 features). Other major functional categories include metabolism of cofactors, vitamins, prosthetic groups, and pigments, respiration, and DNA metabolism. Functional groups related to stress response (76), fatty acids, lipids, and isoprenoids (55), and RNA metabolism (PL152□ and PL155: 55 and 56 features, respectively) are consistently represented in both genomes.

Subsystems associated with membrane transport (PL152□ : 124; PL155: 133), regulation and cell signaling (PL152□ : 58; PL155: 64), and virulence, disease, and defense (PL152□ : 41; PL155: 39) were represented in both genomes. Minor differences were noted in phosphorus metabolism (PL152□ : 27; PL155: 29), iron acquisition (PL152□ : 44; PL155: 42), and nitrogen metabolism (PL152□ : 40; PL155: 43). Subsystem categories with fewer features included motility and chemotaxis (13 in both strains), cell division and cell cycle (7 in both), and secondary metabolism (PL152□ : 8; PL155: 10) (Supplementary Table 4).

The ANI threshold of 95–96%, corresponding to 70% DNA–DNA hybridization (DDH) relatedness, has been recommended and widely adopted for bacterial species delineation (Konstantinidis et al., 2005; Goris et al., 2007; Richter and Rosselló-Móra, 2009; Dobhal et al, 2025). ANI, defining the nucleotide-level genomic similarity of PL152^T^ genome against the strain PL155 was 99.9%; pairwise calculated dDDH values between PL152□ and PL155 was 99.62% suggesting that these belong to the same species in the genus. Pairwise ANI values of PL152^T^ and PL155 against the other Type strains of other species described in the genus *Pectobacterium* ranged from 85.1-93.85% (Table 1; Figure 2). The highest ANI was observed with *P. aroidearum* L6^T^ (93.85%) followed by *P. sinaloense* LFLA-215^T^ (92.73%). In contrast, the lowest ANI was observed with *P. cacticida* CFCC10813 (85.1%) followed by *P. fontis* M022^T^ (88.26%). Pairwise dDDH values between PL152 and PL155 and other *Pectobacterium* species, ranged from 27.7% to 54.3% (Table 1). Consistent with the ANI results, the highest dDDH value was observed with *P. aroidearum* L6(54.3%), followed by *P. sinaloense* LFLA-215^T^ (49.1%). The lowest dDDH similarity was found with *P. cacticida* CFCC10813(27.7%), followed by *P. fontis* M022□ (34.9%). The analyses clearly indicate that strains PL152^T^ and PL155 share dDDH values below the 70% species delineation threshold when compared with other *Pectobacterium* species (Table 1). Therefore, based on the combined ANI and dDDH evidence compared with all recognized species within the genus *Pectobacterium*, we propose *Pectobacterium pacificum* sp. nov., with PL152□ designated as the type strain and PL155 representing an additional strain of this novel species.

**Figure 2.**
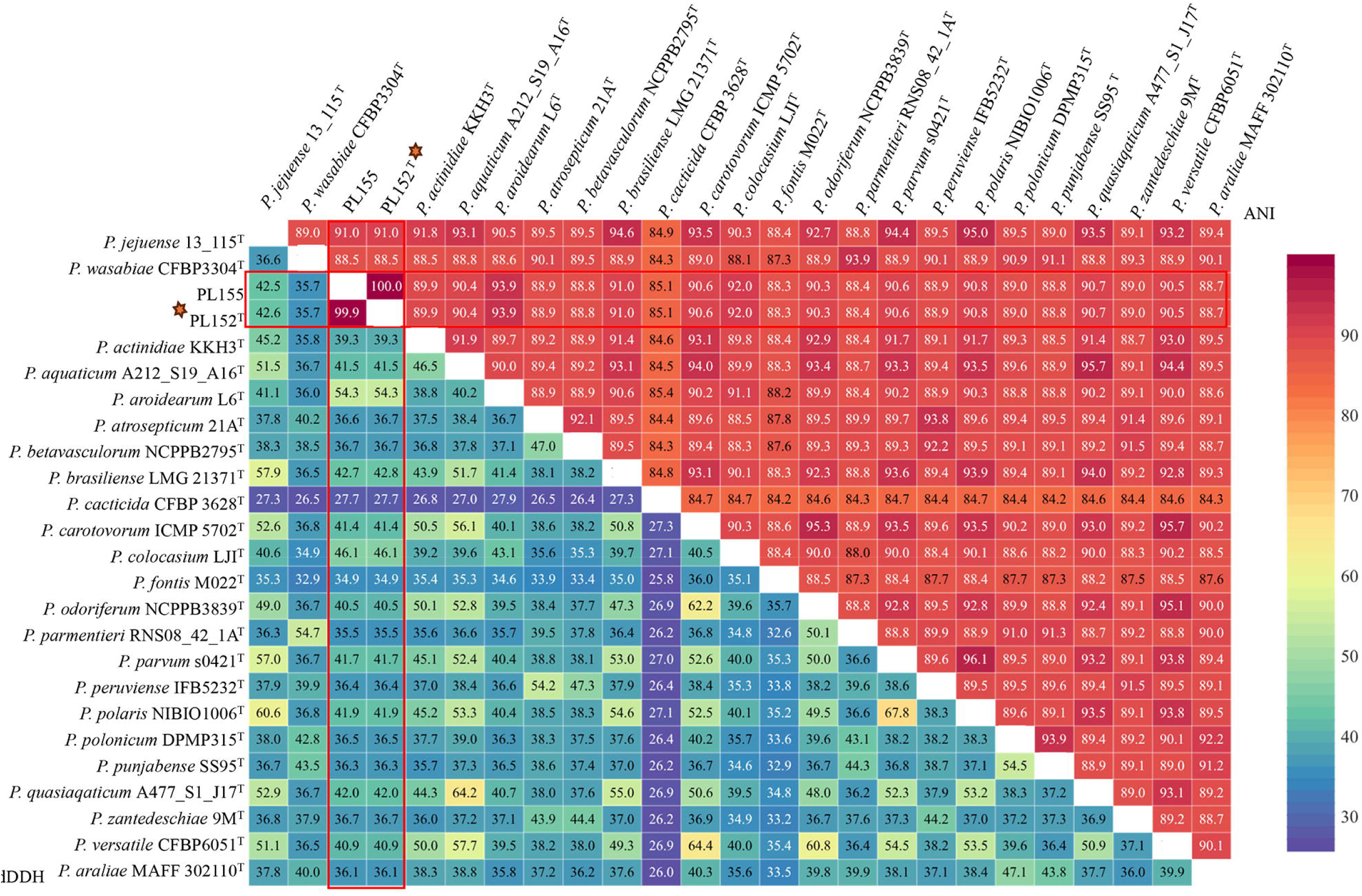
Heatmap showing combined average nucleotide identity (ANI; upper triangle) and digital DNA-DNA hybridization (dDDH; lower triangle) values among the type strains of all described *Pectobacterium* species along with *P. pacificum* sp. nov. strain PL152^T^ and PL155. The species delineation thresholds are 95–96% for ANI and 70% for dDDH. The type strain of the proposed novel species, *P. pacificum* sp. nov. PL152, is indicated by an orange star on the y-axis of the heatmap. The red rectangle indicates the ANI and dDDH values obtained in comparison with other species. *indicates that *Pectobacterium cacticida* has recently been proposed for reclassification as *Alcorniella cacticida* comb. *nov.*; however, the reclassification has not yet been formally approved. The heatmap was generated using Displayr.

**Table 1.** Average nucleotide identity (ANI) and digital DNA-DNA hybridization (dDDH) estimations of *Pectobacterium pacificum* so. nov strains PL152^T^ (=ICMP 24362^T^; LMG 32536 ^T^) and PL155 against all other species within genus *Pectobacterium*. The upper right triangle and lower left triangle represent ANIm and dDDH values, respectively.

| Species | PL152<sup>T</sup> |  | PL155 |  |
| --- | --- | --- | --- | --- |
|  | ANI | DDH | ANI | DDH |
| *P. pacificum* sp. nov. PL152<sup>T</sup> | 100.0 | 100.0 | 99.99 | 99.9 |
| *P. pacificum* sp. nov. PL155 | 99.9 | 99.62 | 100.0 | 100.0 |
| *P. atrosepticum* 21A<sup>T</sup> | 88.89 | 36.7 | 88.89 | 36.6 |
| *P. carotovorum* WPP14<sup>T</sup> | 90.63 | 41.4 | 90.62 | 41.4 |
| *P. actinidiae* KKH3<sup>T</sup> | 89.92 | 39.3 | 89.92 | 39.3 |
| *P. aquaticum* A212_S19_A16<sup>T</sup> | 90.44 | 41.5 | 90.44 | 41.5 |
| *P. araliae* MAFF_302110<sup>T</sup> | 88.73 | 36.1 | 88.72 | 36.1 |
| *P. aroidearum* L6<sup>T</sup> | 93.85 | 54.3 | 93.85 | 54.3 |
| *P. betavasculorum* NCPPB2795<sup>T</sup> | 88.83 | 36.7 | 88.83 | 36.7 |
| *P. brasiliense* 1692<sup>T</sup> | 91.02 | 42.8 | 91.02 | 42.7 |
| *P. cacticida* CFCC10813<sup>T</sup> | 85.1 | 27.7 | 85.1 | 27.7 |
| *P. colocasium* LJ1<sup>T</sup> | 91.96 | 46.1 | 91.96 | 46.1 |
| *P. fontis* M022<sup>T</sup> | 88.26 | 34.9 | 88.26 | 34.9 |
| *P. jejuense* 13_115<sup>T</sup> | 91.04 | 42.6 | 91.04 | 42.5 |
| *P. odoriferum* NCPPB3839<sup>T</sup> | 90.25 | 40.5 | 90.25 | 40.5 |
| *P. parmentieri* RNS08_42_1A<sup>T</sup> | 88.44 | 35.5 | 88.44 | 35.5 |
| *P. parvum* s0421<sup>T</sup> | 90.62 | 41.7 | 90.62 | 41.7 |
| *P. peruviense* IFB5232<sup>T</sup> | 88.88 | 36.4 | 88.88 | 36.4 |
| *P. polaris* NIBIO1006<sup>T</sup> | 90.75 | 41.9 | 90.75 | 41.9 |
| *P. polonicum* DPMP315<sup>T</sup> | 88.95 | 36.5 | 88.95 | 36.5 |
| *P. punjabense* SS95<sup>T</sup> | 88.84 | 36.3 | 88.84 | 36.3 |
| *P. quasiaquaticum* A477_S1_J17<sup>T</sup> | 90.72 | 42.0 | 90.72 | 42.0 |
| *P. versatile* CFBP6051<sup>T</sup> | 90.46 | 40.9 | 90.46 | 40.9 |
| *P. wasabiae* CFBP3304<sup>T</sup> | 88.54 | 35.7 | 88.54 | 35.7 |
| *P. zantedeschiae* 9M<sup>T</sup> | 89.0 | 36.7 | 89.0 | 36.7 |
| *P. sinaloense* LFLA-215<sup>T</sup> | 92.73 | 49.1 | 92.75 | 49.1 |

### 16S rRNA, MLSA and whole geneome phylogenies

The *dnaA* gene is primarily used for the identification of *Pectobacterium* and *Dickeya* species, and it has proven to be an efficient marker for species-level differentiation (Schneider et al, 2011; Dobhal et al, 2020). The further identification was confirmed using the three housekeeping genes (*dnaA*, *gapA*, and *gyrB*). The *gapA* gene has proven to be a reliable marker for the identification and phylogenetic resolution of *Pectobacterium* and *Dickeya* species, enabling discrimination at the genus, species, and subspecies levels (Cigna et al, 2017; Hugouvieux-Cotte-Pattat et al., 2023). The *gyrB* gene has also been widely used for phylogenetic analyses and is considered an efficient marker for bacterial taxonomy (Mainello-Land et al, 2024; Watanabe et al, 2001). The NCBI Blast outcomes confirmed that PL152^T^ and PL155 belong to the genus *Pectobacterium*. Sequence comparison of the *dnaA*, *gyrB*, and *gapA* genes showed that strains PL152 and PL155 shared 96.01%, 95.24%, and 97.7% sequence identity, respectively, with *P. colocasium*. In comparison, the corresponding *dnaA*, *gyrB*, and *gapA* sequences exhibited 95.35%, 98.33%, and 96.2% identity, respectively, with *P. aroidearum*. These results placed PL152 and PL155 close to *P. colocasium* and *P. aroidearum* based on individual housekeeping-gene sequence comparisons. Phylogenetic analysis based on the concatenated *dnaA*, *gyrB*, and *gapA* gene sequences grouped PL152 and PL155 together in a distinct lineage, separate from the recognized *Pectobacterium* species included in the analysis (data not shown).Analysis of the 16S rRNA revealed that PL152^T^ and PL155 clustered with member of genus *Pectobacterium*, confirming their placement within the genus *Pectobacterium* (Supplementary Figure 1).

In the resulting tree, PL152 and PL155 clustered together on short branches, indicating a high level of relatedness and low genetic divergence between the two strains. The PL152 and PL155 strains were positioned near closely related *Pectobacterium* species, particularly *P. aroidearum*, *P. colocasium*, and *P. sinaloense*, but remained separated from the reference strains representing currently described species. This placement suggests that PL152 and PL155 are related to neighboring *Pectobacterium* taxa but form a distinct lineage. Because the 16S rRNA gene is highly conserved and has limited resolution for differentiating closely related species, this analysis was interpreted as preliminary phylogenetic evidence. Together with ANI and dDDH values below accepted species demarcation thresholds, the phylogenetic placement supports the proposal that PL152 and PL155 represent a novel *Pectobacterium* species.

Multi-locus sequence analysis (MLSA) based on nine-housekeeping genes was used to assess the phylogenetic placement of PL152□ and PL155 within the genus *Pectobacterium.* The concatenated dataset comprised *rpoS* (993 bp), *rpoA* (989 bp), *recA* (1,073 bp), *mdh* (931 bp), *gyrB* (736 bp), *gapA* (881 bp), *dnaN* (908 bp), *dnaK* (1,908 bp), and *dnaA* (889 bp), resulting in a total concatenated sequence length of 9,308 bp. The MLSA phylogeny showed that PL152 and PL155 formed a distinct and well-supported clade within the genus *Pectobacterium* (Figure 3). This clade was separated from currently described *Pectobacterium* species, supporting the taxonomic distinctiveness of these strains. PL152□ and PL155 were placed near *P. sinaloense* LFLA-215 and *P. colocasium* LJ1□, forming a neighboring clade to *P. aroidearum* strains L6□, MY2, KC20, and CFBP8737. This placement suggests that *Pectobacterium pacificum* sp. nov. shares a recent common ancestor with these closely related taxa but remains phylogenetically distinct. The strong bootstrap support for the PL152□ -PL155 clade further supports their classification as representatives of a novel species within the genus *Pectobacterium*.

**Figure 3.**
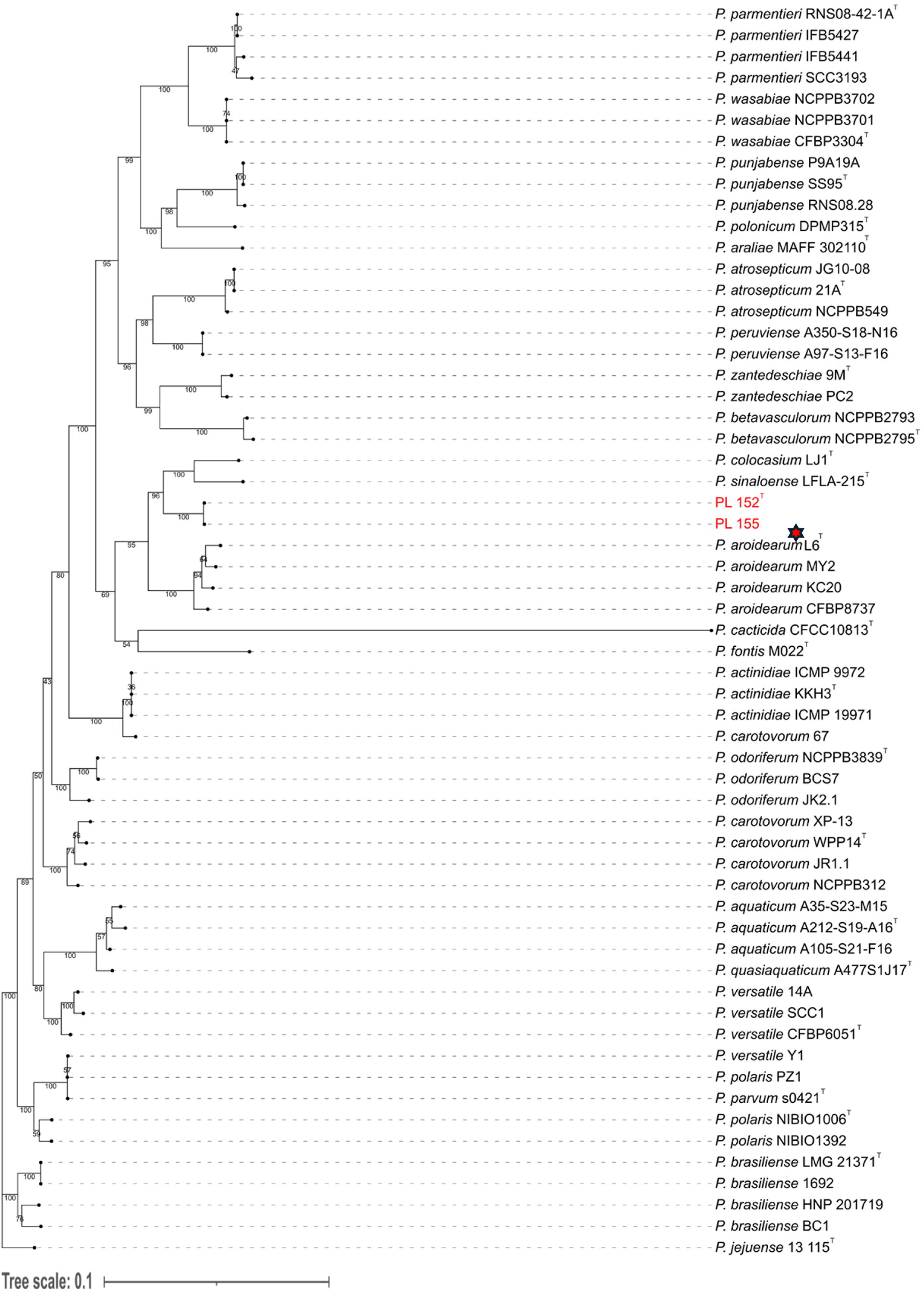
Multi-Locus Sequence Analysis (MLSA) of 57 strains, comprising *Pectobacterium pacificum* (PL152^T^; ICMP 24362^T^; LMG 32536 ^T^ and PL155) and other known *Pectobacterium* sp. was performed using nine housekeeping genes (*dnaA*, *gapA*, *gyrB*, *recA*, *dnaN*, *rpoS*, *mdh*, *rpoA* and *dnaK*). Phylogenetic analysis was performed with Geneious Tree Builder software using MLSA generated in Geneious Prime using RAxML v8.2.11 (GTR+GAMMA) with 1,000 bootstrap replicates. *Pectobacterium pacificum* sp. nov. formed a novel clade.

To further evaluate the taxonomic placement of the proposed novel species, whole-genome-based phylogenetic analysis was performed using the genomes of PL152□ and PL155 together with 24 type-strain genomes representing described species within the genus *Pectobacterium*. In the resulting phylogeny, PL152□ and PL155 clustered together and formed a distinct clade within the genus. This clade was positioned close to *P. aroidearum* L6 and near the clade containing *P. sinaloense* LFLA-215 and *P. colocasium* LJ1 but remained clearly separated from these neighboring taxa. The independent placement of PL152 and PL155 supports their phylogenetic distinctiveness from the most closely related described *Pectobacterium* species (Figure 4). The Type Strain Genome Server TYGS; https://tygs.dsmz.de/) analysis further supported the distinct taxanomic positions of strains PL152□ and PL155. Both PL152□ and PL155 were assigned by TYGS as representing a potential new species, which is consistent with the phylogenetic, ANI, and dDDH analyses. Together, these results support the designation of PL152□ and PL155 as strains of a novel species within the genus *Pectobacterium*, for which the name *Pectobacterium pacificum* sp. nov. is proposed.

**Figure 4.**
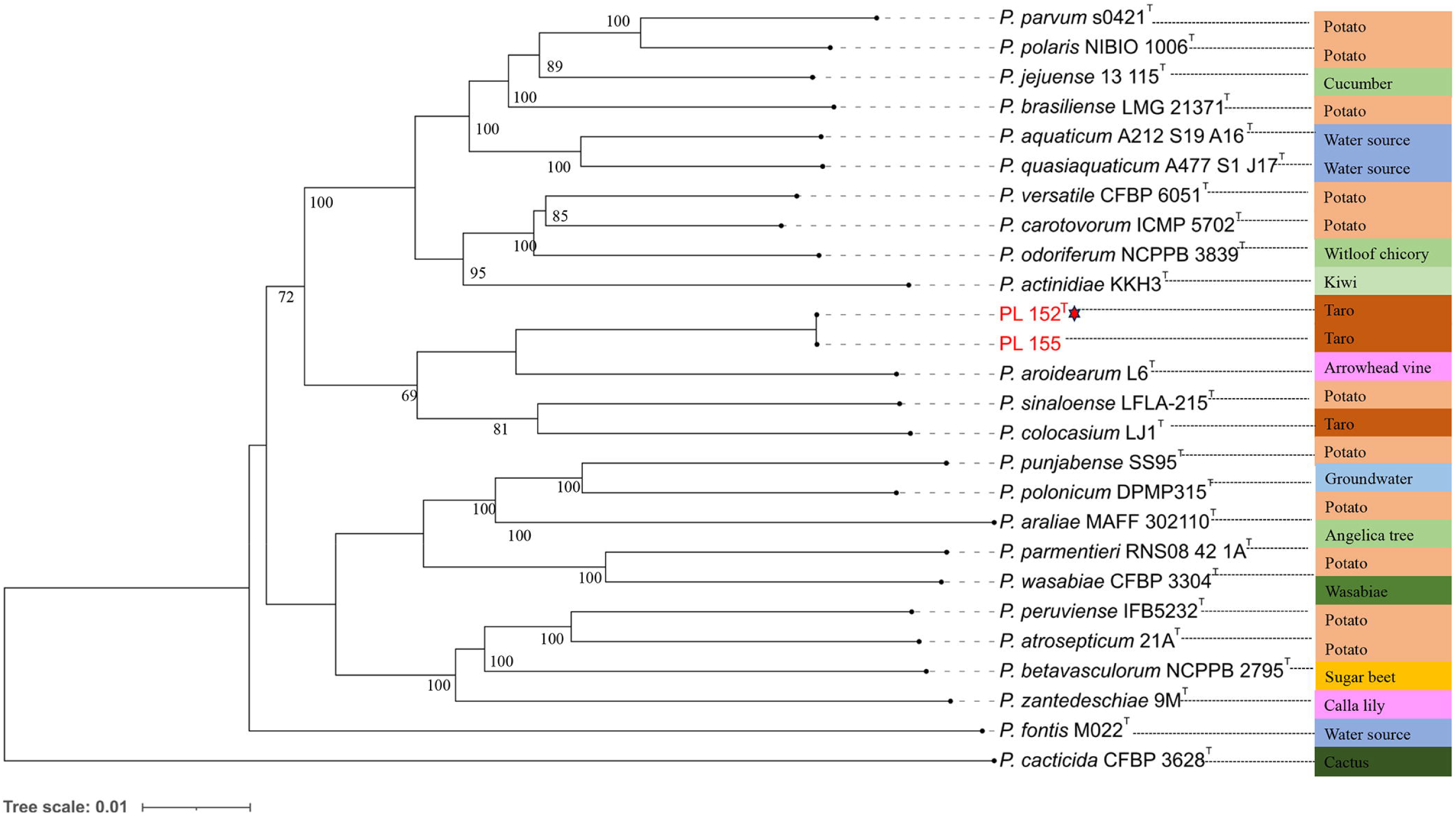
Whole-genome-based phylogenetic tree of the genus *Pectobacterium* showing realtionship between the *P. pacificum* sp. nov. and other species described in the genus. The tree was constucted using Type Strain Genome Server (TYGS) of all *Pectobacterium* species. *Pectobacterium pacificum* formed a distinct clade. The tree was inferred with FastME 2.1.6.1 using the GBDP approach, with distances calculated from genome sequences via the TYGS. Branch lengths are scaled in terms of GBDP distance formula d5. The numbers above the branches are GBDP pseudo-bootstrap support values >60% from 100 replications. The tree was rooted at the midpoint. The Newick file generated by the Type Strain Genome Server (TYGS) was uploaded to the Interactive Tree of Life (iTOL) version 7.0 for visualization and annotation of the phylogenetic tree. Bootstrap analysis was performed using 100 replicates.

### Genomic Analysis

Members of the genus *Pectobacterium* are well known for their ability to cause soft rot and related disease symptoms through a combination of PCWDEs, secretion systems, motility, and host-colonization factors (Arif et al, 2022). The PCWDEs including pectate lyases (Pel), polygalacturonases (Peh), cellulases (Cel), and proteases (Prt) which macerate plant tissues by degrading structural components (Toth et al., 2003; Pérombelon, 2002; Arizala and Arif, 2019). The genomes of PL152 and PL155 were examined for gene clusters associated with pathogenicity, including PCWDEs, secretion systems, chemotaxis, flagellar biosynthesis, and type IV pilus formation. Motility-associated genes were identified in both PL152□ and PL155 genomes (**Supplementary Table 5**). The detected genes included components of the chemotaxis system, flagellar biosynthesis and regulation, flagellar motor function, and type IV pilus biogenesis. Chemotaxis-related genes included *cheA*, *cheW*, *cheR*, *cheB*, *cheY*, *cheZ*, *cheV*, *cheM*, and *cheD*. Genes associated with flagellar assembly and motility, including *flh*, *flg*, *fli*, *motA*, *motB*, and *ycgR*, were also present in both strains. In addition, both genomes contained type IV pilus-associated genes, including *pilL-pilW*, *ppD/pilA*, *hofB/pilB*, and *hofC/pilC*. These findings indicate that both strains possess conserved genetic determinants associated with chemotaxis, flagellum-mediated motility, and pilus-associated surface movement. The genomes of PL152□ and PL155 showed a similar set of PCWDE-associated genes (**Supplementary Table 6**). Several genes involved in pectin breakdown were detected in both genomes, including pectate lyases, pectin lyase, pectin acetylesterases, pectin methylesterases, polygalacturonases, oligogalacturonide lyase, and rhamnogalacturonate lyase. Genes associated with cellulose degradation, including glycoside hydrolase and glucanase-related genes, were also present. In addition, both strains contained multiple protease-associated genes, including metalloproteases, serine proteases, zinc-dependent proteases, aminopeptidases, and related hydrolases.The presence of these enzyme classes suggests that PL152□ and PL155 have the genetic capacity to degrade major components of the plant cell wall. This is consistent with the typical soft rot biology of *Pectobacterium*, where pectin-degrading enzymes play an important role in tissue maceration and disease development (Toth et al., 2006). The bacterial genomes were analyzed for the different gene clusters which were associated with the pathogenesis. The type I secretion system (T1SS) in *Pectobacterium* is a three-component protein complex (prtDEF): an ATP-binding cassette (ABC) transporter (PrtD), a membrane fusion protein (PrtE), and an outer membrane protein of the TolC family (PrtF). The prtDEF operon is responsible for the secretion of metalloproteases (e.g., PrtW), hydrolases, toxins, adhesins, and hemophores from the bacterial cytoplasm to the external environment (Arizala and Arif, 2019; Delepelaire et al., 2004). PL152□ and PL155 strains consisted of prtDEF operon and the cluster involved in the adhesin and proteases secretion were found to be present in both strains (Supplementary Table 7). The T2SS is the most important secretion system, and also known as Out system, allowing secretion of PCWDEs to the extracellular environment (Toth et al., 2006; Arizala and Arif, 2019). The *Out* gene cluster was identified in the genomes of PL152□ and PL155 screened in both genomes, revealing a 17,740 bp region encompassing 17 genes from *gspO* to *gspS*, (Supplementary Table 8; Figure 5). A large T3SS-associated *hrp/hrc* gene cluster of 33 genes of 35,728 bp were identified in both genomes, indicating that these strains possess genetic components related to effector delivery and host interaction (Figure 5).

**Figure 5:**
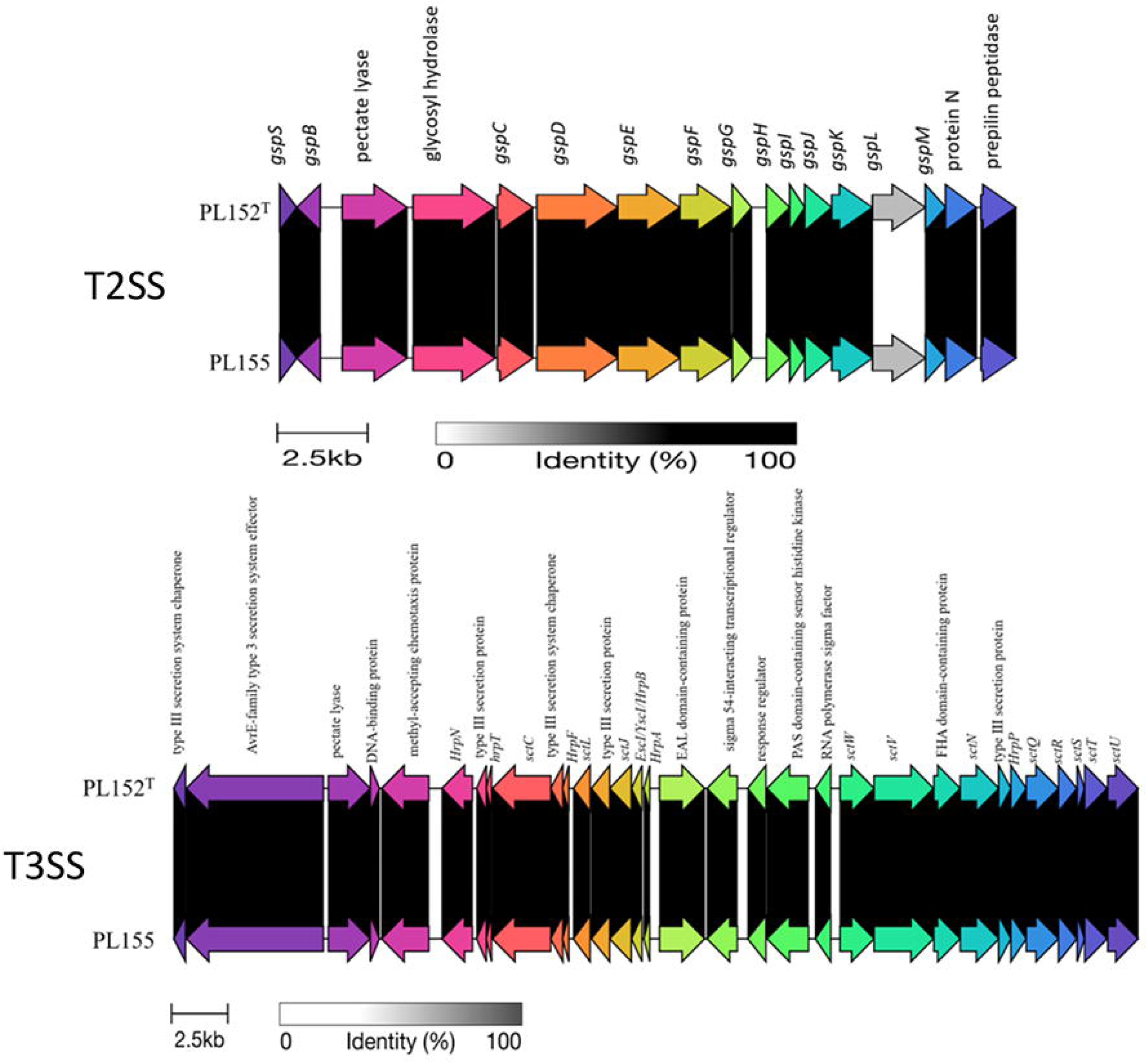
Genetic organization of Type II secretion system (T2SS) and Type III secretion system (T3SS) gene clusters in PL152^T^ and PL155. Comparison of T2SS/T3SS gene clusters between PL152^T^ and PL155. Arrows indicate coding sequence and their orientation. Gene color distinguishes individual genes, and grey shaded regions represent nucleotide identity between two genomes, with darker shading corresponding to higher sequence similarity. The images were generated using clinker and clustermap.js (<u>Gilchrist and Chooi 2021</u>).

In plant-pathogenic bacteria, the T3SS is commonly involved in the translocation of effector proteins into host cells and may contribute to suppression or modulation of plant defense responses (Kim et al., 2009; Chuang et al, 2025). Although soft rot development in *Pectobacterium* is primarily driven by PCWDEs and the T2SS, the presence of T3SS-associated genes suggests that strains PL152□ and PL155 may also possess additional virulence-related traits that contribute to host colonization and plant-pathogen interactions (Figure 5). Putative type V two-partner secretion system associated genes were identified in both genomes. These included genes encoding ShlB/FhaC/HecB family secretion or activation proteins together with hemagglutinin repeat-containing proteins and two-partner secretion domain-containing proteins. The presence of these paired transporter proteins suggests that both strains possess T5SS-associated adhesin systems, which may contribute to surface attachment, bacterial aggregation, and host colonization. An additional locus containing a ShlB/FhaC/HecB family protein, a toxin-activating lysine-acyltransferase, and a hemagglutinin repeat-containing protein was also detected in both genomes (Supplementary Table 9). These findings indicate that the strains carry conserved T5SS-related loci; however, their specific role in adhesion, secretion, or pathogenicity requires further functional validation. The genomes were screened for type VI secretion system (T6SS)-associated genes, and a putative T6SS gene cluster was identified. This region contained several core structural and accessory components, including *paar*, *vgrG*, *hcp*, *vasL*, *tssM/icmF/vasK*, *tssA/impA*, *vasI*, *vasH*, *clpV/tssH*, *tssL, tssK/impJ/vasE*, *tssJ/vasD*, *tagH/impI/vasC*,*tssG/impH/vasB*, *tssF/impG/vasA*, *tssE*, *tssC/impC/vipB*, and *tssB/impB/vipA*.

The presence of *hcp*, *vgrG*, and *paar* suggests that this genomic region contain genes associated with the T6SS secretion apparatus and effector delivery module (Supplementary Table 10). In addition, the detection of sheath, baseplate, ATPase, and regulatory components supports the presence of a T6SS-associated region in this genome. Overall, the detected T6SS-associated genes suggest that both strain may possess genetic potential for interbacterial competition, environmental fitness, and interactions within plant-associated microbial communities, although functional activity would require experimental validation.

Comparative analysis using Panaroo identified gene content changes in PL152□ and PL155. A total of 448 genes were identified as gained, while 4,781 genes were lost relative to closely related strains (Supplementary Table 11-14). This pattern indicated substantial genome remodeling, with gene loss being the dominant process. Func□ ional categorization of gained and lost genes revealed distinct patterns. Gained genes were mainly associated with transport systems, defense mechanisms, mobile genetic elements, and regulatory functions, along with a large number of hypothetical proteins. Together, the observed patterns of gene loss and selective gene acquisition suggest potential functional specialization in PL152 and PL155.

AMRFinderPlus 4.0.23 identified genes (*arsA, arsB, arsC, arsD and asrR*) related to arsenate resistance in both genomes, suggesting that these strains may have the ability to cope with arsenic-related stress. These genes could help the organism survive in environments containing arsenic or other metal stressors. In many bacteria, resistance to arsenic is typically controlled by *ars* operon genes that regulate, transform, and remove toxic arsenic compounds from the cell (Rosen, 2002; Mukhopadhyay et al., 2002). Therefore, the detection of these genes points to a possible adaptive advantage under such conditions. Similar arsenic resistance determinants, including *arsC*, *arsB*, and *arsR*, have been reported in several *Pectobacterium* species, supporting their role in environmental adaptation (Arizala and Arif, 2019; Valdez-Lopez et al., 2026). However, further laboratory experiments would be necessary to confirm whether these both strains can actually tolerate arsenic exposure. AntiSMASH (V8.0.4) analysis identified a predicted biosynthetic gene cluster classified as a mixed NRP-metallophore, NRPS, phosphonate, and T1PKS-associated cluster. The cluster showed high similarity to the known amonabactin P750 biosynthetic gene cluster (from *Aeromonas hydrophila* subsp. *hydrophila* ATCC 7966), suggesting that PL152^T^ and PL155 strains may have the genetic potential to produce a siderophore or metallophore like secondary metabolite. This type of cluster may be associated with metal acquisition, and environmental adaptation. *Pectobacterium* species has also been reported for the production of antimicrobial compounds which includes carotovoricin or carocins with toxic effect against closely realted bacteria (Chuang et al 2007; Chan et al 2009; Roh et al 2010; Arizala and Arif, 2019). We also found *Carocin D* gene present in both genomes of *P. pacificum* sp. nov. The BAGEL4 analysis revealed three putative bacteriocin/RiPP (ribosomally synthesized and post-translationally modified peptides) associated areas of interest (AOIs), corresponding to Carocin_D, microcin, and bottromycin-like regions. The Carocin D like region aligns with previously reported carocin type bacteriocins in *Pectobacterium* species, supporting its likely relevance in this species (Shyntum et al., 2019). In contrast, the microcin and bottromycin like AOIs are interpreted as predictions derived from genome mining rather than evidence of confirmed compound production. Microcins are small antibacterial peptides typically associated with Gram-negative bacteria, particularly members of the Enterobacteriaceae, whereas bottromycins are RiPP type antibacterial compounds predominantly described in *Streptomyces* species (Cole et al., 2022; Vior et al 2020). The identification of these predicted regions suggests that these strains may possess genetic potential for antimicrobial compound production and interbacterial competition; however, further experimental validation is required to confirm their expression and biological activity. DefenseFinder detected 48 genes spanning 17 prokaryotic antiviral defense systems in both genomes (Supplementary Table 15), indicating that these genomes contain multiple mechanism that may withstand phage infection. The presence of these defense associated systems suggests a potential role in genome stability and survival in competitive environments (Valdez-Lopez et al., 2026). The CRISPR-CAS system has been reported to provide the adpative immunity in bacteria and three types of CRIPSR-CAS syatems have been reported in *Pectobacterium* species (Cady et al., 2012; Makarova et al., 2015; Arizala and Arif., 2019; Li et al., 2019). The CRISPR-CASFinder predicted CRISPR-cas systems in both PL152^T^ and PL155. Two types of CRISPR-Cas system Class 1 subtypeIII A and subtype 1F were predicted by CRISPR-CASFinder in both PL152^T^ and PL155. In PL152^T^ and PL155 the CAS-TypeIIIA locus consisted of *cas10*, *csm2*, *csm3*, *cas1*, and *cas2* genes, whereas TypeIF contained *cas1*, *cas3-cas2*, *csy1*, *csy2*, *csy3*, and *cas6*. High confidence CRISPR arrays which includes 5, 10, 6, 17, and 47 spacers were located near these cas clusters which may contribute to the genome stability and immunity (Supplementary Table 16).

### Phenotypic, Biochemical, enzymatic, pathogenecity and antibiotic assays

Colonies on nutrient agar supplemented with 0.4% dextrose were smooth, moist consistency and ceramic white on solid SOB media (2% tryptone, 0.5% yeast extract, 10 mM NaCl, 2.5 mM KCl, 10 mM MgSO_4_, 1.5% agar supplemented with 2% glycerol) after 2 days at 28°C. Electron microscopy showed that *P. pacificum* sp. nov. strains PL152 and PL155 had rod-shaped cells with peritrichous flagella (Figure 6). Cells of PL152 and PL155 measured approximately 2.64 ± 0.27 μm and 2.66 ± 0.27 μm in length, respectively, and approximately 1.00 ± 0.08 μm in width. These measurements represent the mean of 20 observations obtained from three independent replicates. The observed cell morphology and flagellation pattern are consistent with traits reported for motile members of the soft rot *Pectobacteriaceae* (Rungnapha et al., 2008). PL152 and PL155 exhibited motility as indicated with the swimming and swarming assays which were positive for both strains.

**Figure 6.**
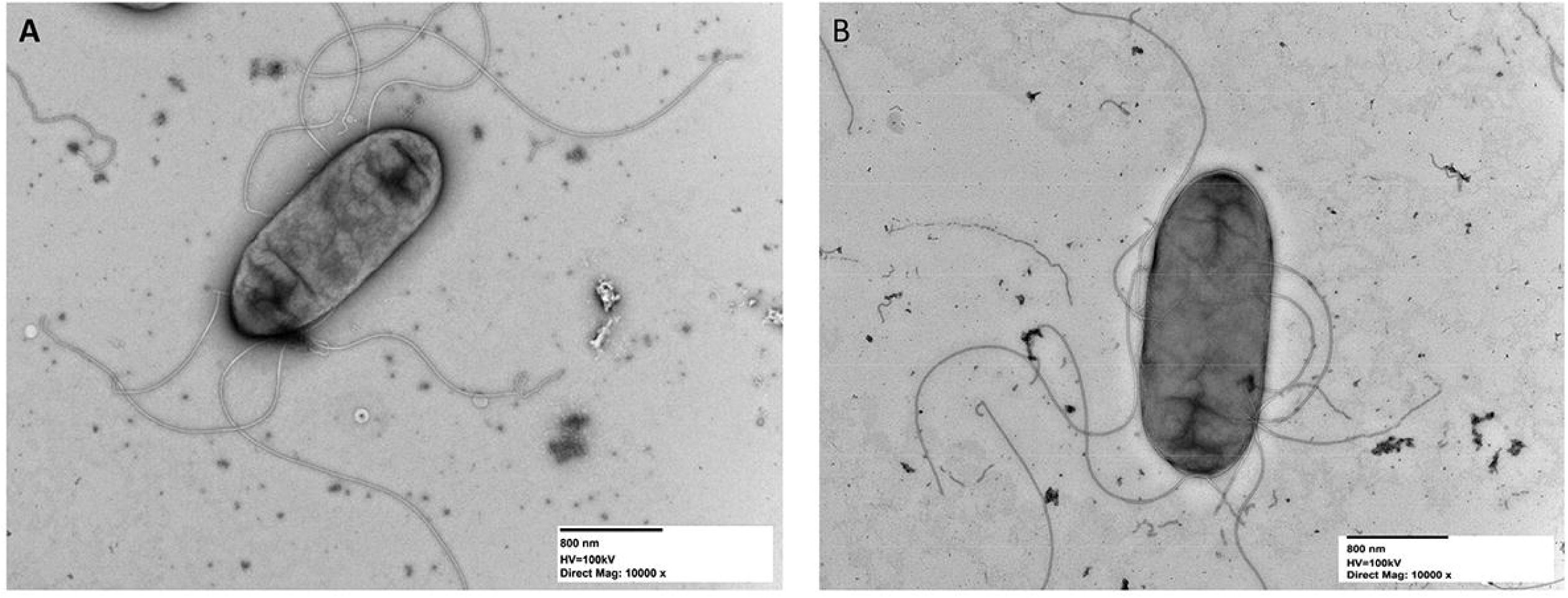
Transmission electron micrograph of *P. pacificum* sp. nov. strain (A) PL152^T^ and (B) PL155. A minimum of ten images were captured for the analyzed strain across three biological replicates. A representative image is shown. The scale bar indicates 800 nm.

Phenotypic and biochemical characterization of PL152□ and PL155 was performed using Biolog GEN III MicroPlates, which include 94 phenotypic assays comprising 71 carbon-source utilization tests and 23 chemical-sensitivity assays. Both strains showed positive utilization of α-D-glucose, pectin, α-D-lactose, D-mannose, D-mannitol, D-galacturonic acid, methyl pyruvate (which was negative in *P. sinaloense* LFLA-125□, *P. colocasium* LJ1□, and weakly utilized by *P. aroidearum* LJ2□), D-fructose, L-galactonic acid lactone, β-methyl-D-glucoside, D-galactose, myo-inositol, D-gluconic acid, D-cellobiose, D-salicin, glycerol, L-aspartic acid, citric acid, gentiobiose, N-acetyl-D-glucosamine, D-glucose-6-phosphate, sucrose, D-fructose-6-phosphate, mucic acid, L-rhamnose, D-aspartic acid, L-malic acid, acetic acid, D-serine, L-serine, D-saccharic acid, and bromo-succinic acid (Table 2), similar to the closely related *P. sinaloense* LFLA-125□, *P. colocasium* LJ1□, and *P. aroidearum* LJ2□, which also utilize many of these substrates (Zhou et al., 2022; Valdez-Lopez et al., 2026). Weak positive reactions were observed for dextrin, D-melibiose, D-trehalose, L-glutamic acid, glucuronamide, acetoacetic acid, and formic acid. Similar to *P. sinaloense* LFLA-125□, *P. colocasium* LJ1□, and *P. aroidearum* LJ2□, neither strain utilized propionic acid but, in contrast, showed a weak reaction for D-serine. In addition, both strains were able to tolerate 1% NaCl, 1% sodium lactate, troleandomycin, lincomycin, vancomycin, pH 6, 4% NaCl, rifamycin SV, guanidine HCl, tetrazolium violet, lithium chloride, niaproof 4, and tetrazolium blue.

**Table 2.** Phenotypic characterization of *Pectobacterium pacificum* PL152^T^ (=ICMP 24362^T^; LMG 32536 ^T^) and PL155 using Biolog GEN III MicroPlate for analysis of 71 carbon source utilization assays and 23 chemical sensitivity assays. Microplates were incubated at 28°C for 24 hours and read using Biolog’s Microbial Identification System software.

| Characteristics (Carbon | PL152<sup>T</sup> | PL155 | Characteristics (Carbon | PL152<sup>T</sup> | PL155 |
| --- | --- | --- | --- | --- | --- |

| Source/Chemical<br>Sensitivity) |  |  | Source/Chemical<br>Sensitivity) |  |  |
| --- | --- | --- | --- | --- | --- |
| D-Raffinose | - | - | Sucrose | + | + |
| α-D-Glucose | + | + | N-Acetyl-β-D-Mannosamine | - | - |
| D-Sorbitol | - | - | L-Fucose | - | - |
| Gelatin | - | - | D-Fructose-6-PO<sub>4</sub> | + | + |
| Pectin | + | + | L-Histidine | - | - |
| p-Hydroxy-Phenylacetic Acid | - | - | Mucic Acid | + | + |
| Tween 40 | - | - | D-Malic Acid | - | - |
| Dextrin | wr | wr | Propionic Acid | - | - |
| α-D-Lactose | + | + | D-Turanose | - | - |
| D-Mannose | + | + | N-Acetyl-D-Galactosamine | - | - |
| D-Mannitol | + | + | L-Rhamnose | + | + |
| Glycyl-L-Proline | - | - | D-Aspartic Acid | + | + |
| D-Galacturonic Acid | + | + | L-Pyroglutamic Acid | - | - |
| Methyl Pyruvate | + | + | Quinic Acid | - | - |
| γ-Amino-Butyric Acid | - | - | L-Malic Acid | + | + |
| D-Maltose | - | - | Acetic Acid | + | + |
| D-Melibiose | wr | wr | Stachyose | - | - |
| D-Fructose | + | + | N-Acetyl Neuraminic Acid | - | - |
| D-Arabitol | - | - | Inosine | - | - |
| L-Alanine | - | - | D-Serine | + | + |
| L-Galactonic Acid Lactone | + | + | L-Serine | + | + |
| D-Lactic Acid Methyl Ester | - | - | D-Saccharic Acid | + | + |
| α-Hydroxy-Butyric Acid | - | - | Bromo-Succinic Acid | + | + |
| D-Trehalose | wr | wr | Formic Acid | wr | wr |
| β-Methyl-D-Glucoside | + | + | 1% NaCl | + | + |
| D-Galactose | + | + | 1% Sodium Lactate | + | + |
| myo-Inositol | + | + | Troleandomycin | + | + |
| L-Arginine | - | - | Lincomycin | + | + |
| D-Gluconic Acid | + | + | Vancomycin | + | + |
| L-Lactic Acid | - | - | Nalidixic Acid | - | - |
| β-Hydroxy-D,L-Butyric Acid | - | - | Aztreonam | - | - |
| D-Cellobiose | + | + | pH 6 | + | + |
| D-Salicin | + | + | 4% NaCl | wr | wr |
| 3-Methyl Glucose | - | - | Fusidic Acid | wr | wr |
| Glycerol | + | + | Rifamycin SV | + | + |
| L-Aspartic Acid | + | + | Guanidine HCl | + | + |
| D-Glucuronic Acid | - | - | Tetrazolium Violet | + | + |
| Citric Acid | + | + | Lithium Chloride | + | + |
| α-Keto-Butyric Acid | - | - | Sodium Butyrate | wr | wr |
| Gentiobiose | + | + | pH 5 | - | - |
| N-Acetyl-D-Glucosamine | + | + | 8% NaCl | - | - |
| D-Fucose | - | - | D-Serine | wr | wr |
| D-Glucose-6-PO4 | + | + | Minocycline | - | - |
| L-Glutamic Acid | wr | wr | Niaproof 4 | + | + |
| Glucuronamide | wr | wr | Tetrazolium Blue | + | + |
| α-Keto-Glutaric Acid | - | - | Potassium Tellurite | - | - |
| Acetoacetic Acid | wr | wr | Sodium Bromate | - | - |

wr-weak reaction; + positive reaction; - negative reaction

Both PL152□ and PL155 did not utilize D-raffinose, D-sorbitol, gelatin, p-hydroxy-phenylacetic acid, Tween 40, glycyl-L-proline, γ-amino-butyric acid, D-maltose, D-arabitol, L-alanine, D-lactic acid methyl ester, α-hydroxy-butyric acid, L-arginine, L-lactic acid, β-hydroxy-D,L-butyric acid, 3-methyl glucose, D-glucuronic acid, α-keto-butyric acid, D-fucose, α-keto-glutaric acid, N-acetyl-β-D-mannosamine, L-fucose, L-histidine, D-malic acid, propionic acid, D-turanose, N-acetyl-D-galactosamine, L-pyroglutamic acid, quinic acid, stachyose, N-acetylneuraminic acid, or inosine (Table 2). The key phenotypic characteristics differentiating PL152□ and PL155 from the closely related species *P. sinaloense* LFLA-125□, *P. colocasium* LJ1□, and *P. aroidearum* LJ2□ are summarized in Supplementary Table 17.

The API 20E system is a commercial biochemical assay used to support the identification of members of the *Enterobacteriaceae* and other non-fastidious Gram-negative rod-shaped bacteria (Dobhal et al., 2025). Tests suitable for identifying the genus and characterizing the species are included. Both *P. pacificum* sp. nov. strains, PL152^T^ and PL155, were positive for indole and acetoin production, citrate utilization, exhibited β-galactosidase activity and negative for gelatinase activity (Table 3). Both strains also fermented D-glucose, D-mannitol, inositol, L-rhamnose, D-sucrose, amygdalin, L-arabinose and negative for D-sorbitol and D-melibiose. However, they were negative for arginine dihydrolase, lysine decarboxylase, ornithine decarboxylase, H2S production, urease and tryptophane deaminase tests. Nitrate reduction to nitrite was also observed for both strains (Table 3).

**Table 3.**
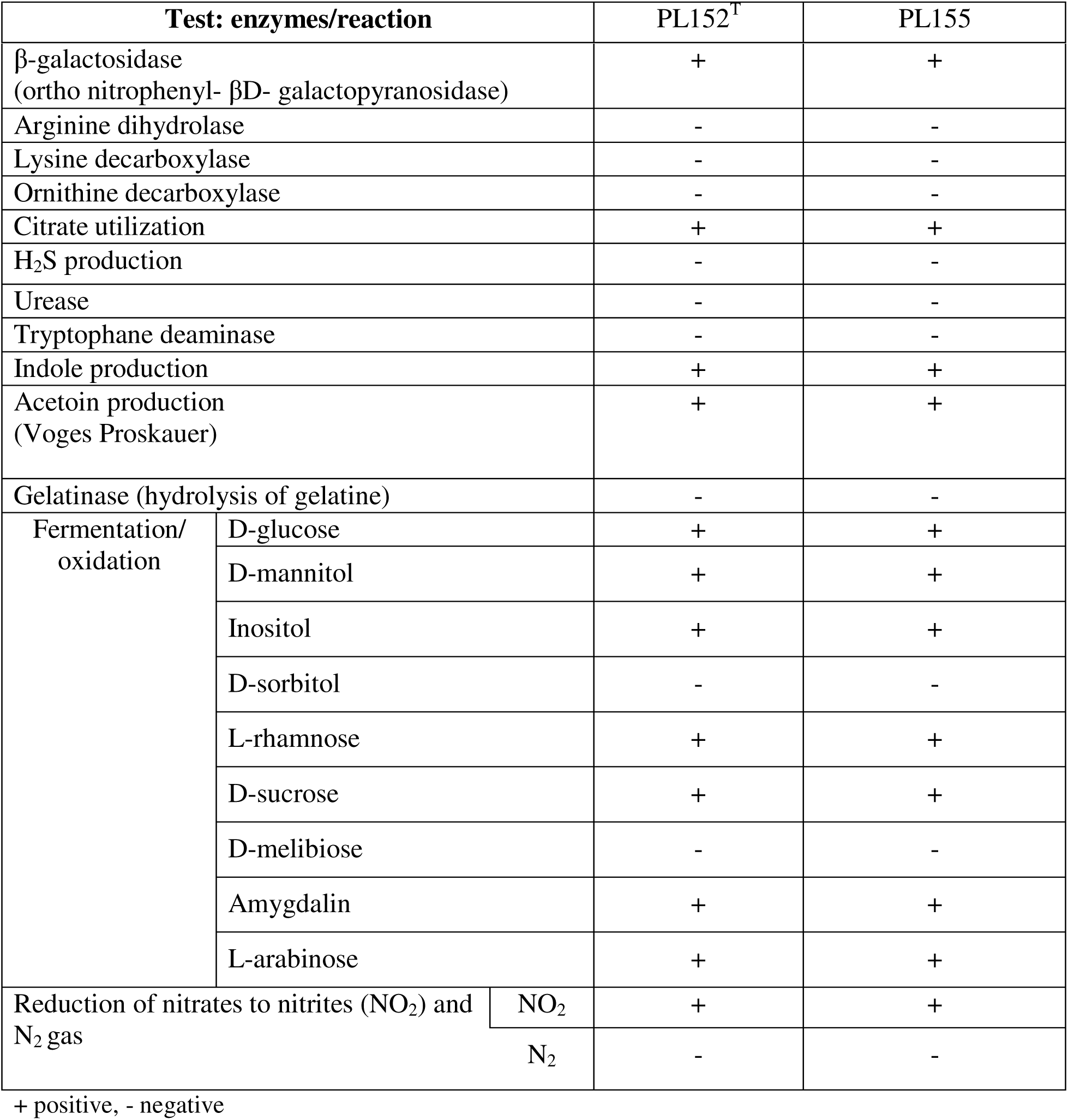
Enzymatic and fermentation analyses of the new species *Pectobacterium pacificum* sp. nov. PL152**^T^** and PL155 based on the API 20E test

The API ZYM test for enzymatic activity revealed positive reactions for alkaline phosphatase, esterase (C4), leucine arylamidase, acid phosphatase, naphthol-AS-BI-phosphohydrolase, α-galactosidase, β-galactosidase, α-glucosidase, and β-glucosidase. Both strains showed negative reactions for esterase lipase (C8), lipase (C14), α-chymotrypsin, β-glucuronidase, N-acetyl-β-glucosaminidase, α-mannosidase, and α-fucosidase. Weak reactions were observed for valine arylamidase, cystine arylamidase, and trypsin (Table 4).

**Table 4.**
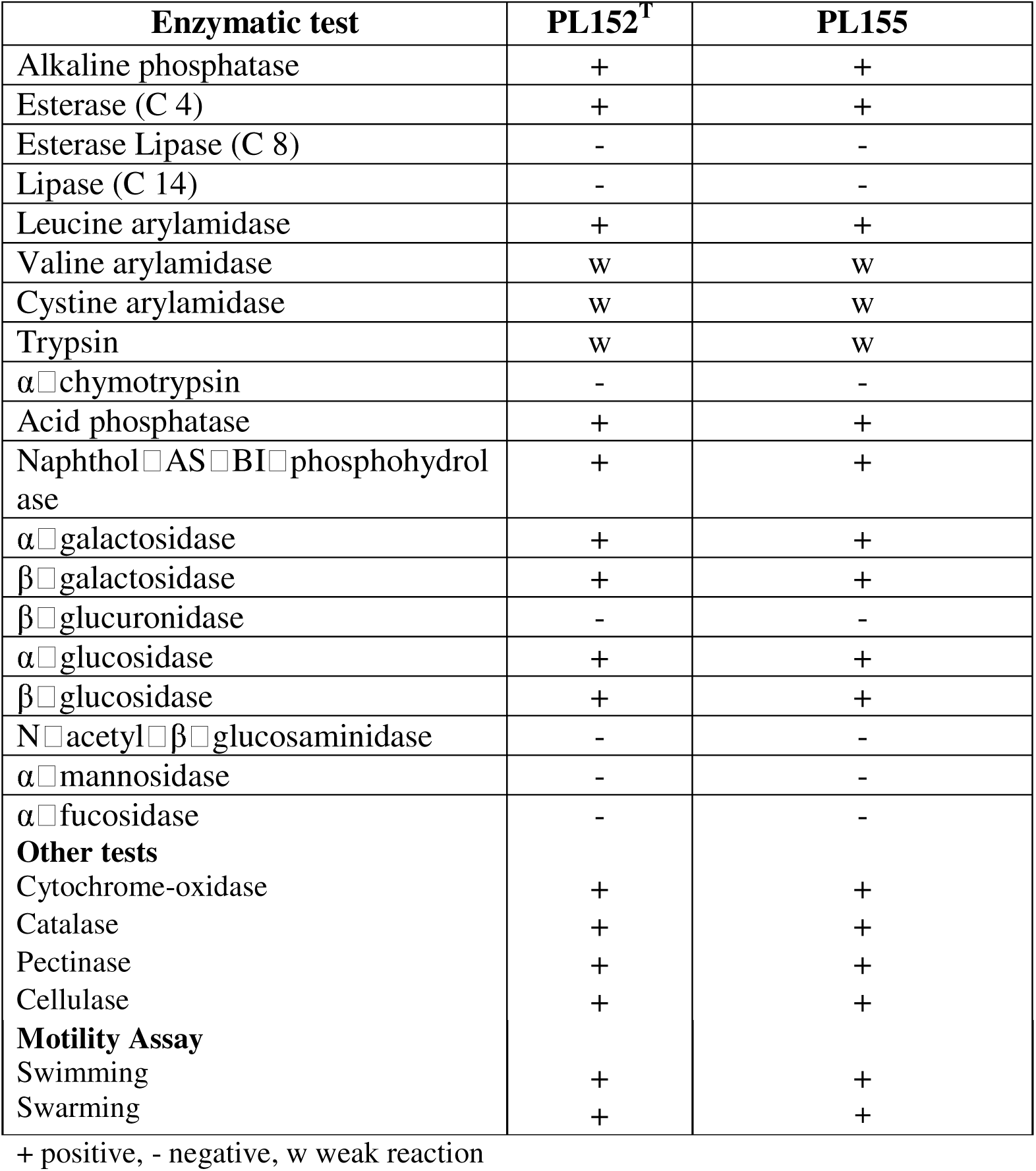
Analyses of enzymatic activities of the new species *Pectobacterium pacificum* sp. nov. PL152**^T^** and PL155 using the API ZYM test strip and other tests.

| Enzymatic test | PL152<sup>T</sup> | PL155 |
| --- | --- | --- |
| Alkaline phosphatase | + | + |
| Esterase (C 4) | + | + |
| Esterase Lipase (C 8) | - | - |
| Lipase (C 14) | - | - |
| Leucine arylamidase | + | + |
| Valine arylamidase | w | w |
| Cystine arylamidase | w | w |
| Trypsin | w | w |
| α-chymotrypsin | - | - |
| Acid phosphatase | + | + |
| Naphthol-AS-BI-phosphohydrolase | + | + |
| α-galactosidase | + | + |
| β-galactosidase | + | + |
| β-glucuronidase | - | - |
| α-glucosidase | + | + |
| β-glucosidase | + | + |
| N-acetyl-β-glucosaminidase | - | - |
| α-mannosidase | - | - |
| α-fucosidase | - | - |
| **Other tests** |  |  |
| Cytochrome-oxidase | + | + |
| Catalase | + | + |
| Pectinase | + | + |
| Cellulase | + | + |
| **Motility Assay** |  |  |
| Swimming | + | + |
| Swarming | + | + |

+ positive, - negative, w weak reaction

Pectate lyases, cellulases, and proteases are essential extracellular enzymes produced by *Dickeya* species, contributing significantly to bacterial pathogenesis (Arif et al, 2022; Boluk et al, 2021; Hugouvieux-Cotte-Pattat et al, 2014). These enzymes facilitate infection by degrading plant cell walls and tissues, thereby enabling pathogen entry, colonization, and spread within the host (Boluk et al, 2021; Hugouvieux-Cotte-Pattat et al, 2014). The role of these PCWDEs is well established in soft rot *Pectobacteriaceae*, with studies confirming their central role in virulence of both *Pectobacterium* and *Dickeya* species across a range of crops, including potato and taro (Alič et al., 2024; Waleron et al., 2019b; Boluk et al, 2021). Both strains, PL152^T^ and PL155, showed positive activity for cytochrome-oxidase, catalase, pectinase, and cellulase, consistent with the enzymatic profiles typical of pathogenic *Pectobacterium* and *Dickeya* isolates (Loc et al, 2022). Both strains, PL152□ and PL155, were able to degrade pectin, as demonstrated by pit formation on crystal violet pectate medium. Pathogenicity assays performed on potato tubers and taro corms showed maceration on both, indicating positive pathogenic activity and confirming the ability of the tested bacterial strains, PL152^T^ and PL155, to infect and degrade plant tissues. Antibiotic sensitivity profiling serves as a valuable phenotypic tool for characterizing *Pectobacterium* and *Dickeya* species (Dobhal et al., 2025; Tan et al, 2022). It also provides insights into their ecological fitness and potential survival strategies across diverse environments. Variability in antibiotic resistance among isolates from different hosts and ecological niches underscores the utility of antibiotic susceptibility tests in species delineation, while also highlighting their importance in understanding ecological adaptation and informing potential treatment strategies (Hossain et al, 2023; Toth et al, 2021). Furthermore, the novel species was sensitive to the seven tested antibiotics namely bacitracin, chloramphenicol, kanamycin, carbeicillin, tetracycline, gentamicin and penicillin. PL152^T^ and PL155 showed different sensitivity to antibiotics by forming inhibition zone diameters of 44-and 51-mm for penicillin (50 mg/ml), 20-and 21.5-mm for kanamycin (50 mg/ml), 28-and 37-mm for tetracycline (40 mg/ml), 46-and 35.5-mm chloramphenicol (50 mg/ml), 32-and 37.5-mm carbenicillin (100 mg/ml), 16-and 21-mm gentamicin (50 mg/ml) and 8-and 12.5-mm bacitracin (50 mg/ml). The characteristics of the new proposed species *P. pacificum* sp. nov. are listed in the protologue description in Table 5.

**Table 5.**
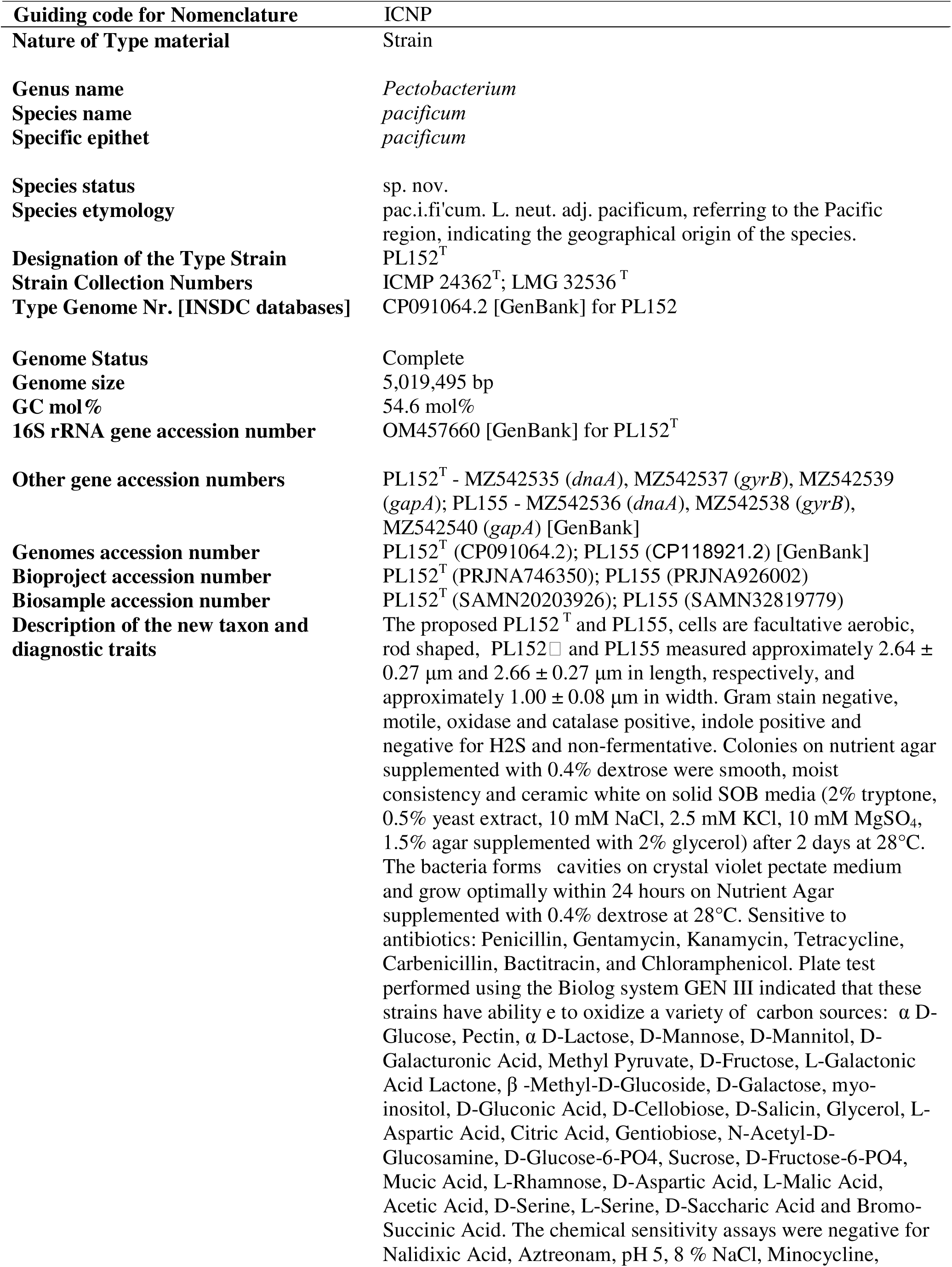
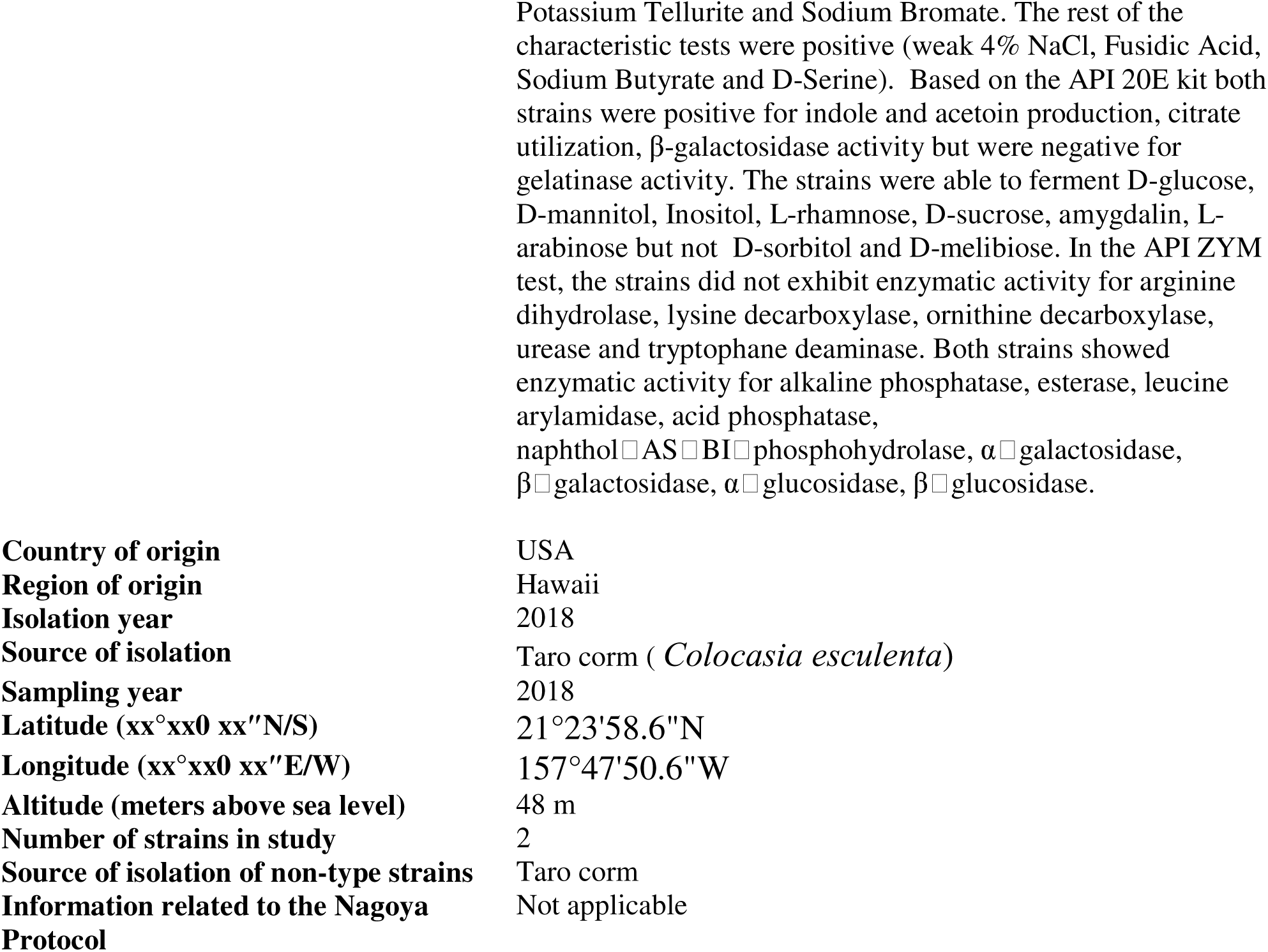
Protologue description of *Pectobacterium pacificum* sp. Nov.

|  |  |
| --- | --- |
| Guiding code for Nomenclature | ICNP |
| **Nature of Type material** | Strain |
| **Genus name** | *Pectobacterium* |
| **Species name** | *pacificum* |
| **Specific epithet** | *pacificum* |
| **Species status** | sp. nov. |
| **Species etymology** | pac.i.fi'cum. L. neut. adj. pacificum, referring to the Pacific region, indicating the geographical origin of the species. |
| **Designation of the Type Strain** | PL152<sup>T</sup> |
| **Strain Collection Numbers** | ICMP 24362<sup>T</sup>; LMG 32536<sup>T</sup> |
| **Type Genome Nr. [INSDC databases]** | CP091064.2 [GenBank] for PL152 |
| **Genome Status** | Complete |
| **Genome size** | 5,019,495 bp |
| **GC mol%** | 54.6 mol% |
| **16S rRNA gene accession number** | OM457660 [GenBank] for PL152<sup>T</sup> |
| **Other gene accession numbers** | PL152<sup>T</sup> - MZ542535 (*dnaA*), MZ542537 (*gyrB*), MZ542539 (*gapA*); PL155 - MZ542536 (*dnaA*), MZ542538 (*gyrB*), MZ542540 (*gapA*) [GenBank] |
| **Genomes accession number** | PL152<sup>T</sup> (CP091064.2); PL155 (CP118921.2) [GenBank] |
| **Bioproject accession number** | PL152<sup>T</sup> (PRJNA746350); PL155 (PRJNA926002) |
| **Biosample accession number** | PL152<sup>T</sup> (SAMN20203926); PL155 (SAMN32819779) |
| **Description of the new taxon and diagnostic traits** | The proposed PL152<sup>T</sup> and PL155, cells are facultative aerobic, rod shaped, PL152<sup>T</sup> and PL155 measured approximately 2.64 ± 0.27 µm and 2.66 ± 0.27 µm in length, respectively, and approximately 1.00 ± 0.08 µm in width. Gram stain negative, motile, oxidase and catalase positive, indole positive and negative for H<sub>2</sub>S and non-fermentative. Colonies on nutrient agar supplemented with 0.4% dextrose were smooth, moist consistency and ceramic white on solid SOB media (2% tryptone, 0.5% yeast extract, 10 mM NaCl, 2.5 mM KCl, 10 mM MgSO<sub>4</sub>, 1.5% agar supplemented with 2% glycerol) after 2 days at 28°C. The bacteria forms cavities on crystal violet pectate medium and grow optimally within 24 hours on Nutrient Agar supplemented with 0.4% dextrose at 28°C. Sensitive to antibiotics: Penicillin, Gentamycin, Kanamycin, Tetracycline, Carbenicillin, Bactitracin, and Chloramphenicol. Plate test performed using the Biolog system GEN III indicated that these strains have ability to oxidize a variety of carbon sources: α D-Glucose, Pectin, α D-Lactose, D-Mannose, D-Mannitol, D-Galacturonic Acid, Methyl Pyruvate, D-Fructose, L-Galactonic Acid Lactone, β -Methyl-D-Glucoside, D-Galactose, myo-inositol, D-Gluconic Acid, D-Cellobiose, D-Salicin, Glycerol, L-Aspartic Acid, Citric Acid, Gentioibiose, N-Acetyl-D-Glucosamine, D-Glucose-6-PO<sub>4</sub>, Sucrose, D-Fructose-6-PO<sub>4</sub>, Mucic Acid, L-Rhamnose, D-Aspartic Acid, L-Malic Acid, Acetic Acid, D-Serine, L-Serine, D-Saccharic Acid and Bromo-Succinic Acid. The chemical sensitivity assays were negative for Nalidixic Acid, Aztreonam, pH 5, 8 % NaCl, Minocycline, |
| **Country of origin** | USA |
| **Region of origin** | Hawaii |
| **Isolation year** | 2018 |
| **Source of isolation** | Taro corm (*Colocasia esculenta*) |
| **Sampling year** | 2018 |
| **Latitude (xx°xx' xx"N/S)** | 21°23'58.6"N |
| **Longitude (xx°xx' xx"E/W)** | 157°47'50.6"W |
| **Altitude (meters above sea level)** | 48 m |
| **Number of strains in study** | 2 |
| **Source of isolation of non-type strains** | Taro corm |
| **Information related to the Nagoya Protocol** | Not applicable |

### Conclusions

In conclusion, comprehensive polyphasic analyses including 16S rRNA sequencing, multilocus sequence analysis of nine housekeeping genes, whole-genome sequencing, ANI, and dDDH, clearly distinguish strains PL152 and PL155 from all currently recognized species within the genus *Pectobacterium*. These strains, isolated from taro corms in Hawai‘i, form a distinct and well-supported clade within the genus. Phenotypic, biochemical, and pathogenicity assays further support their uniqueness. Both strains produced extracellular enzymes such as pectate lyase and cellulase, tested positive for virulence on taro corms and potato tubers, and displayed distinct antibiotic sensitivity patterns. Based on the genomic and phenotypic evidence, we propose the establishment of *Pectobacterium pacificum* sp. nov., with PL152 (=ICMP 24362; LMG 32536) as the type strain. This taxonomic delineation advances the understanding of species diversity and ecological adaptation within the genus *Pectobacterium*.

## CRediT authorship contribution statement

**Mohammad Arif**: conceptualization, writing, review and editing, analysis, project administration, funding acquisition, investigation, supervision, resources. **Komal**: writing, data curation, analysis, review and editing. **Diksha Klair**: writing partial preliminary draft, methodology. **Dario Arizala:** review and editing, methodology. **Aarya Mishra: analysis and review. Gamze Boluk:** review and editing, preliminary investigation, methodology. **Małgorzata Waleron**: review and editing. **Anne M. Alvarez:** review and editing**. Shefali Dobhal**: conceptualization, methodology, analysis, data curation, writing, review, editing,, and funding acquisition.

## Acknowledgements

The authors sincerely thank Dr. Bevan Weir and Ms. Megan Petterson from ICMP, New Zealand, as well as Ms. Els Vercoutere and Ms. Claudine Vereecke from the BCCM/LMG Bacteria Collection, for their valuable support in facilitating the registration of strains in the respective culture collections.

## Data Availability

The nucleotide sequence of the 16S rRNA gene for PL152 and PL155 were deposited in the NCBI GenBank database under accession number OM457660 and PV652463, respectively. The genome of the type strain PL152, is available under accession number CP091064.2, with the associated BioProject PRJNA746350 and BioSample SAMN20203926. The other strain PL155 is deposited under the following accession numbers: CP118921.2, with the associated BioProject PRJNA926002 and BioSample SAMN32819779.

## SUPPLEMENTARY MATERIALS

Figure S1. The 16S rRNA tree was generated using Type Strain Genome Server and visualized and annotated using Interactive Tree of Life (iToL) version 7.0. for all type strains within *Pectobacterium*: *P. colocasium* LJ1^T^, *P. actinidiae* KKH3^T^, *P. carotovorum* WPP14^T^, *P. aroidearum* L6^T^, *P. brasiliense* 1692^T^, *P. polaris* NIBIO1006^T^, *P. parvum* s0421^T^, *P. odoriferum* NCPPB3839^T^, *P. versatile* CFBP6051^T^, *P. aquaticum* A212-S19-A16^T^, *P. zantedeschiae* 9M^T^, *P. fontis* M022^T^, *P. polonicum* DPMP315^T^, *P. punjabense* SS95^T^, *P. parmentieri* RNS08-42-1A^T^, *P. wasabiae* CFBP3304^T^, *P. atrosepticum* 21A^T^, *P. peruviense* IFB5232^T^, *P. betavasculorum* NCPPB2795^T^, *P. quasiaquaticum* A477-S1-J17^T^, *P. cacticida* CFCC10813^T^, *P. jejuense* 13-115^T^, *P. araliae* MAFF 302110^T^ and *P. sinaloense* LFLA-215^T^. *Pectobacterium pacificum* sp. nov. strain PL152^T^ (ICMP 24362^T^; LMG 32536 ^T^) and PL155 rooted among *Pectobacterium* species. The evolutionary history was inferred using the Maximum Likelihood method with a bootstrap of 1,000 replicates. The analysis involved 26 strains.

**Supplemental Table 1.** List of type strains of known *Pectobacterium* species, retrieved from the NCBI GenBank database and used for the 16S rRNA analysis.

**Supplemental Table 2.** List of 57 bacterial strains of *Pectobacterium* species retrieved from NCBI GenBank database for multi-locus sequence analysis.

**Supplementary Table 3.** Genome assembly and quality statistics of the *Pectobacterium* strains PL152^T^ and PL155.

**Supplementary Table 4.** The Subsystem Annotation summary from the RAST web server categorizes features into 28 distinct groups.

**Supplementary Table 5.** The flagellar, chemotaxis, and twiching motility gene clusters present and their locus tag in both PL152□ and PL155 genomes of *P. pacificum* sp. nov.

**Supplementary Table 6.** The Plant cell wall degrading enzymes (PCWDEs) associated genes detected in both PL152□ and PL155 genomes of *P. pacificum* sp. nov.

**Supplementary Table 7.** Type 1 Secretion System (T1SS) cluster genes with the locus tag detected in both PL152□ and PL155 genomes of *P. pacificum* sp. nov.

**Supplementary Table 8:** Type II Secretion System genes detected in both PL152□ and PL155 genomes of *P. pacificum* sp. nov.

**Supplementary Table 9.** Type V Secretion System (T5SS) cluster genes with the locus tag detected in both PL152□ and PL155 genomes of *P. pacificum* sp. nov.

**Supplementary Table 10.** Type VI Secretion system genes and effectors detected in both PL152□ and PL155 genomes of *P. pacificum* sp. nov.

**Supplementary Table 11.** Genes uniquely gained in *Pectobacterium pacificum* sp. nov. relative to closely related *Pectobacterium* species.

**Supplementary Table 12.** Functional distribution of genes gained in *Pectobacterium pacificum* sp. nov.

**Supplementary Table 13.** Genes lost from *P. pacificum* sp. nov. compared with closely related *Pectobacterium* species.

**Supplementary Table 14.** Functional distribution of genes lost in *Pectobacterium pacificum* sp. nov.

**Supplementary Table 15**. DefenseFinder detected anti-phage defense system identified in the genomes of PL152□ and PL155.

**Supplementary Table 16.** Detailed description of CRISPR and *CAS* gene clusters found in genomes of PL152□ and PL155.

**Supplementary Table 17.** Key phenotypic characteristics differentiating PL152□ and PL155 from the closely related species *P. sinaloense* LFLA-125□, *P. colocasium* LJ1□, and *P. aroidearum* LJ2□. Data included for the comparison were included from Zhou et al. 2022 and Valdez-Lopez et al., 2026.

