## Supplementary figures and images for "Description of *Pectobacterium pacificum* sp. nov. isolated from *Colocasia esculenta*"

### Figure S1

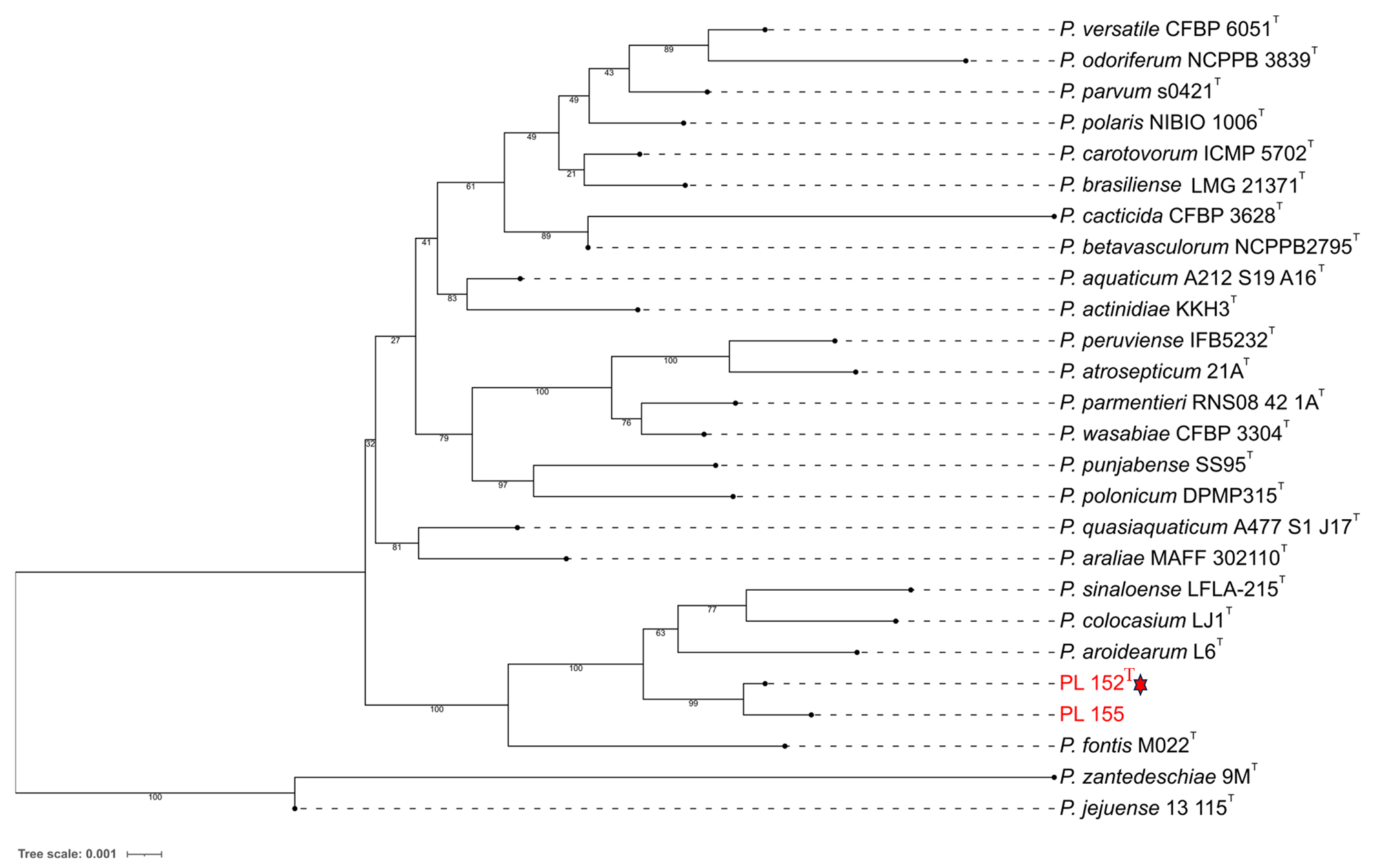
